# Prevalent regulation of GATA2/3 and MSX2 on endogenous retrovirus-derived regulatory elements in human trophoblast stem cells

**DOI:** 10.1101/2022.08.02.502490

**Authors:** Cui Du, Jing Jiang, Yuzhuo Li, Miao Yu, Jian Jin, Shuai Chen, Hairui Fan, Todd S. Macfarlan, Bin Cao, Ming-an Sun

## Abstract

The placenta is an organ with extraordinary phenotypic diversity in eutherian mammals. Recent evidence suggests that numerous human placental enhancers are evolved from lineage-specific insertions of endogenous retroviruses (ERVs), yet the transcription factors (TFs) underlying their regulation remain largely elusive. Here, by first focusing on MER41, a primate-specific ERV family previously linked to placenta and innate immunity, we uncover the binding motifs of multiple crucial trophoblast TFs (GATA2/3, MSX2, GRHL2) in addition to innate immunity TFs STAT1 and IRF1. Integration of ChIP-Seq data confirms the binding of GATA2/3, MSX2 and their related factors on the majority of MER41-derived enhancers in human trophoblast stem cells (TSCs). Notably, MER41-derived enhancers that are constitutively active in human TSCs are distinct from those activated upon interferon stimulation, which is determined by the binding of relevant TFs and their sub-family compositions. We further demonstrate that GATA2/3 and MSX2 have prevalent binding on numerous other ERV families – indicating their broad impact on ERV-derived enhancers. Functionally, the derepression of many syncytiotrophoblast genes after disruption of MSX2 is likely to be mediated by regulatory elements derived from ERVs – suggesting ERVs are also important for mediating transcriptional repression. Overall, this study characterized the prevalent regulation of GATA2/3, MSX2 and their co-factors on ERV-derived regulatory elements in human TSCs and provided mechanistic insights into the importance of ERVs in human trophoblast regulatory network.

## Introduction

Endogenous retroviruses (ERVs) make up ∼8% of the human genome. ERVs originate from germline infection by exogenous retroviruses, which can then be vertically inherited and expanded in the host during evolution (Stoye 2012; Johnson 2019). The intact structure of ERVs includes the internal protein-coding genes (eg. *gag, pol and env*) flanked by Long Terminal Repeats (LTRs) at both ends, yet the internal genes frequently got lost via LTR recombination resulting in a high proportion of “solo-LTRs” left in the genome (Johnson 2019). ERVs have long been ignored as “junk DNA” - partly because most of them are epigenetically repressed (Sharif et al. 2016; Deniz et al. 2019; Wolf et al. 2020), yet accumulating evidence suggests that specific ERV families could be activated during embryonic development, tumorigenesis or immune response (Macfarlan et al. 2012; Chuong et al. 2016; Schoenfelder et al. 2018; Todd et al. 2019; Ito et al. 2020). Importantly, many ERV families are lineage-specific and rich of transcription factor binding sites (TFBSs) in their LTRs, therefore can be co-opted as cis-regulatory elements to boost the genetic novelty of the host (Chuong et al. 2016; Chuong et al. 2017; Senft and Macfarlan 2021; Sun et al. 2021; Buttler and Chuong 2022).

The placenta is a temporary organ crucial for nutrient/waste exchange, hormone secretion and maternal-fetus immune tolerance during pregnancy (Maltepe and Fisher 2015; Ander et al. 2019). Even though shared by all eutherian mammals, placentae are extraordinarily diversified in their shape, structure and cellular composition (Ramsey et al. 1976; Mossman 1987; Hemberger et al. 2020). At the molecular level, numerous genes have altered expression patterns in human placenta relative to rodents or even other primates (Rosenkrantz et al. 2021; Sun et al. 2021). Correspondingly, the enhancers of placenta are the least conserved across species compared with other tissues (Sun et al. 2021), with ERVs playing crucial roles for lineage-specific enhancer evolution. For example, comparison between mouse and rat trophoblast stem cells (TSCs) indicates that the mouse-specific RLTR13D5 family creates hundreds of enhancers co-bound by the core trophoblast transcription factors (TFs) Cdx2, Eomes and Elf5 (Chuong et al. 2013). The functional importance of ERVs in primate placentae has also been analyzed recently. For example, one primate-specific THE1B element creates an enhancer driving *CRH* expression to influence gestation length (Dunn-Fletcher et al. 2018). Recent transcriptomic and epigenomic comparisons for human, macaque and mouse placentae further identified dozens of ERV families that are significantly enriched in human placental enhancers (Sun et al. 2021).

Despite the well-recognized importance of ERVs in the placenta, the transcription factors that regulate ERV activation in the human placenta remain largely obscure. Previously we demonstrated that the primate-specific MER41 is amongst the top most enriched ERV families in human placental enhancers (Sun et al. 2021). MER41 has six sub-families including A/B/C/D/E/G (Kojima 2018), that have been shown to play important roles in innate immunity (Buttler and Chuong 2022). Multiple studies suggest that MER41 elements facilitate primate-specific innate immunity evolution by creating hundreds of interferon (IFN)-stimulated cis-elements bound by STAT1 and IRF1 (Schmid and Bucher 2010; Chuong et al. 2016; Raviram et al. 2018). This raises the question of which TFs mediate the activation of MER41-associated enhancers in placenta. We previously identified Serum response factor (SRF) as one TF that binds dozens of MER41-associated enhancers (Sun et al. 2021), including one adjacent to *FBN2*, a human-placenta-expressed gene crucial for cell invasion (Yu et al. 2020). However, SRF only binds a relatively small percentage (∼6%) of MER41-associated enhancers, indicating the existence of additional upstream TFs. Uncovering the regulators of ERV-associated enhancers will be crucial for an in-depth understanding of the gene regulatory network in human placenta.

In this study, we aim at identifying the TFs that potentially regulate the cis-regulatory elements associated with MER41 transposons and other ERV families in human trophoblast stem cells. By in-depth characterization of the genome-wide binding and regulatory function of the identified candidate TFs, we expect this study will improve our understanding about the regulatory mechanism and function of ERV-associated enhancers in human placenta.

## Results

### Tissue-specific activation of MER41-associated enhancers in human placenta and trophoblast cells

The primate-specific MER41 family creates numerous lineage-specific enhancers in human placenta (Sun et al. 2021). To better characterize MER41-associated enhancers (hereafter abbreviated as MER41-enhancer), we examined the epigenetically-annotated enhancers (ie. H3K27ac peaks that are more than 500 bp away from any TSSs) in different human tissues and trophoblast cells. Re-analysis of the ChIP-Seq data of H3K27ac from ENCODE project (**Table S1**) demonstrates that multiple subfamilies of MER41 (A/B/C/D) are specifically enriched in the enhancers of placenta (as well as the chorion, a fetally derived extraembryonic membrane) relative to other human tissues (**Fig. 1A; Supplemental Fig. S1, S2**). Furthermore, genes adjacent to MER41-enhancers have increased expression in human placenta relative to other tissues and mouse placenta (**Fig. 1B; Supplemental Fig. S3**), and some enriched GO terms (eg. endocytosis, chordae embryonic development) for MER41-enhancers are placenta-related (**Supplemental Fig. S4**). Further inspection suggests that the genes with increased expression in human placenta relative to mouse are most significantly enriched surrounding enhancers derived from MER41B/D, followed by moderate enriched surrounding MER41C-enhancers (**Supplemental Fig. S5**).

**Figure 1.**
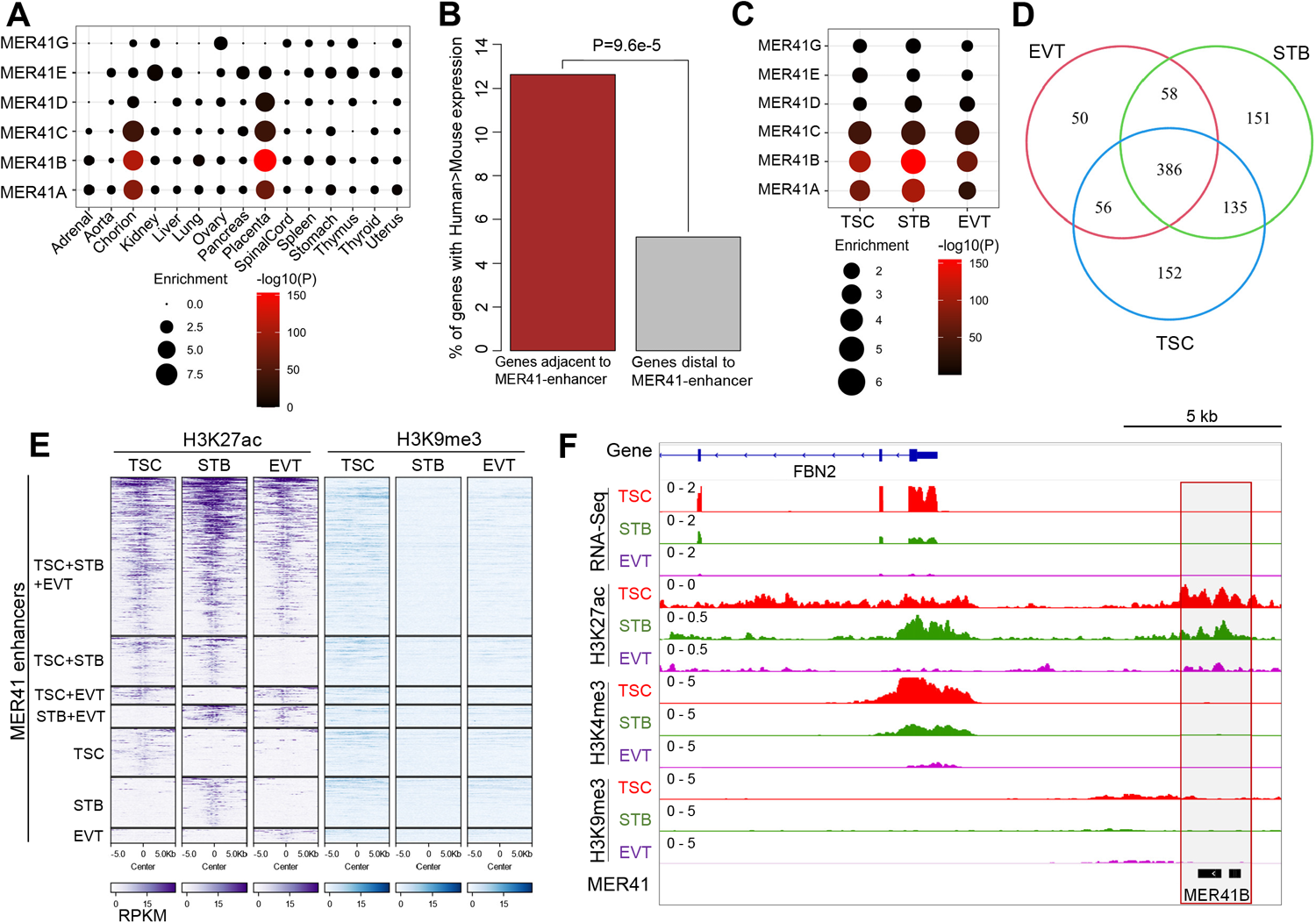
Characterization of MER41-enhancers in different human tissues and trophoblast cells. **(A)** Enrichment of MER41 subfamilies in the epigenetically-annotated enhancers for different human tissues. Enhancers were annotated as H3K27ac peaks that are more than 500 away from any TSSs. **(B)** The bar plot shows that, relative to distal genes, higher proportion of MER41-enhancer adjacent genes have increased expression in human placenta relative to mouse. MER41-enhancer adjacent genes are defined as those with their TSSs within 50 KB from any MER41-enhancers. P-value calculated using Fisher’s Exact Test is indicated. **(C)** Enrichment of MER41 subfamilies in the enhancers annotated for human TSC, STB and EVT. **(D)** Overlapping of the MER41-enhancers annotated for human TSC, STB and EVT. **(E)** Heatmap shows the H3K27ac and H3K9me3 intensity for the shared or cell-specific MER41-enhancers for different human trophoblast cells. **(F)** IGV tracks showing the transcriptomic and epigenomic profiles surrounding the representative MER41-enhancer adjacent to *FBN2*.

Through integration of published epigenomic data (Okae et al. 2018), we further annotated the enhancers for human TSC, syncytiotrophoblast (STB) and extravillous trophoblast (EVT) cells, which show substantial differences among these cell types (**Supplemental Fig. S6**). MER41 elements are enriched in the enhancers for all three trophoblast cell types (**Fig. 1C**). Notably, hundreds of MER41-enhancers are specifically activated in one or two trophoblast cell types (**Fig. 1D,E; Table S2**), and their chromatin state correlates with the transcription of at least some adjacent genes, such as *FBN2* and *C1QTNF6* (**Fig. 1F, Supplemental Fig. S7**). Together, these results confirm the tissue-specific activation of MER41-associated enhancers in human placenta and indicate their potential regulatory effects on some adjacent genes.

### Identification of GATA2/3 and MSX2 as putative regulators of MER41-enhancers in human TSCs

To screen for candidate TFs that bind MER41-enhancers, we first determined the motif composition of MER41B consensus sequence and one representative MER41-enhancer adjacent to *FBN2*. Apart from several TFs (ie. STAT1, IRF1, SRF) already known to bind MER41 elements (Chuong et al. 2016; Sun et al. 2021), the binding motifs for dozens of other TFs are predicted (**Fig. 2A,B)**. Importantly, four of them, including GATA2/3, MSX2 and GRHL2, have placenta-enriched expression (**Fig. 2C; Supplemental Fig. S8**). Particularly, GATA2/3 are key pioneer TFs for human and mouse TSC specification (Home et al. 2017; Krendl et al. 2017; Paul et al. 2017) and MSX2 is essential in restraining STB gene expression in human TSCs (Hornbachner et al. 2021). Even though GRHL2 has not been analyzed in human placenta, it is known to regulate trophoblast branching morphogenesis in mice (Walentin et al. 2015). Inspection of other subfamilies including MER41A/C/D also gave similar results (**Supplemental Fig. S9**), and direct comparison of the MER41-enhancers against MER41-elements that lacking the H3K27ac mark uncovered the enrichment of the motifs for GATA3 and GRHL2 (**Supplemental Fig. S10**), but not MSX2 which is assumed for transcriptional repression. Therefore, motif prediction and transcriptomic analysis uncovered GATA2/3, MSX2 and GRHL2 as candidate trophoblast TFs that bind MER41-enhancers in human placenta.

**Figure 2.**
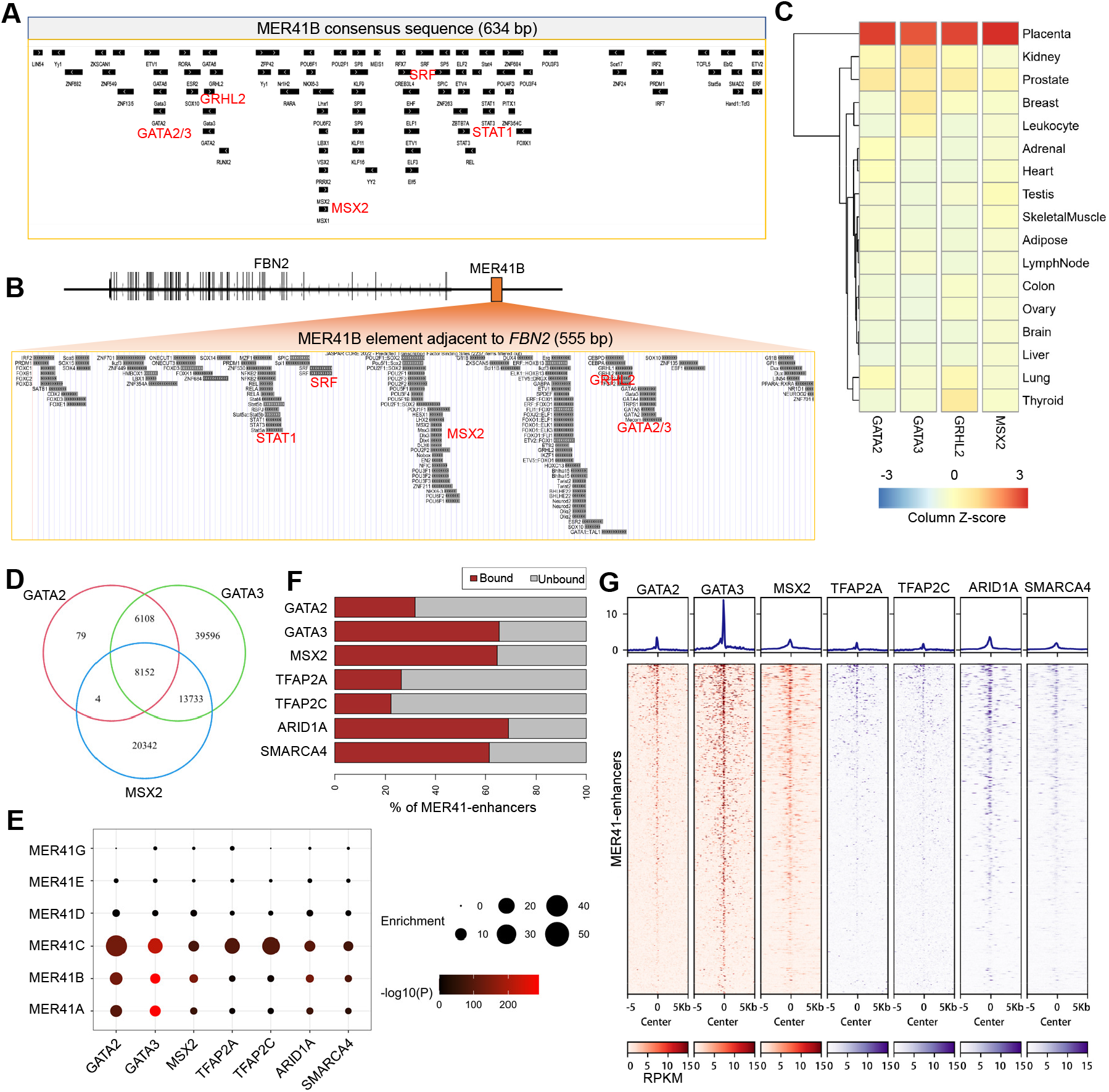
Integrative analysis identified GATA2/3 and MSX2 as candidate regulators of MER41-enhancers in human TSCs. **(A,B)** Motif configurations of MER41B consensus (A) and one representative MER41-enhancer adjacent to *FBN2* (B) The motifs for several known (STAT1, SRF) and novel (GATA2/3, MSX2, GRHL2) TFs of MER41 are labelled in red color. **(C)** Expression profiles of the four TFs, including *GATA2/3, MSX2* and *GRHL2*, that have placenta-enriched expression. The color gradient indicates the column Z-score based on normalized TPM values. **(D)** Venn diagram showing the peak overlapping of GATA2/3 and MSX2. **(E)** Enrichment of MER41 subfamilies in the peaks for GATA2/3, MSX2 and co-factors. **(F)** Bar plots demonstrate the percentages of MER41-enhancers that are bound by GATA2/3, MSX2 and co-factors. (**G**) Heatmaps demonstrate the prevalent binding of GATA2/3, MSX2 and co-factors on MER41-enhancers in human TSCs.

Recent studies have profiled the genome-wide binding of GATA2/3 (together with TFAP2A/C which are collectively coined as the “trophectoderm four”) and MSX2 (together with its co-factors ARID1A and SMARCA4) in human TSCs by using ChIP-Seq (Krendl et al. 2017; Hornbachner et al. 2021). Through re-analysis of these data, we demonstrate that GATA2/3 and MSX2 have substantially overlapped binding (**Fig. 2D**), and all these TFs and co-factors have enriched binding on MER41-enhancers in human TSCs (**Fig. 2E; Table S3**). Notably, the majority (>60%) of 634 MER41-enhancers annotated for human TSC are bound by GATA3 and MSX2 (and its co-factors ARID1A and SMARCA4), as indicated by both overlapping analysis and heatmap visualization (**Fig. 2F,G**). The remaining TFs also have widespread binding on MER41-enhancers, even though of lower frequency (**Fig. 2F,G**). These results were further confirmed by manual inspection of a few representative loci, such as *FBN2*, and *C1QTNF6* (**Supplemental Fig. S11**). Together, our integrative analyses identified several crucial trophoblast TFs, including GATA2/3 and MSX2, as putative regulators of MER41-enhancers in human TSCs.

### Direct comparison of MER41-enhancers that are constitutively-active or IFN-stimulated in human TSCs

Intrigued by previous findings that MER41 elements facilitate the evolution of lineage-specific enhancers that are either constitutively activated in normal placenta (Sun et al. 2021) or interferon-stimulated in different human cells (Chuong et al. 2016), we directly compared between placenta- and immune-related MER41-enhancers using human TSC as model. For this purpose, we performed transcriptomic and epigenomic (H3K27ac) profiling of human TSCs with or without interferon gamma (IFNG) stimulation (**Supplemental Table S1**). As expected, IFNG stimulates hundreds of genes in human TSCs which are highly associated with immune response (**Supplemental Fig. S12**). We next determined thousands of IFN-stimulated enhancers (**Supplemental Fig. S13**) and further examined their overlap with MER41 subfamilies. While MER41A/B/C are enriched with similar fold within constitutively-active enhancers, MER41B is enriched within IFN-stimulated enhancers to a higher degree (**Fig. 3A**). Impressively, the constitutively-active enhancers – in contrast to IFN-stimulated ones – are indeed unresponsive to IFN-stimulation (**Fig. 3B,C; Table S4**), and closer inspection of representative loci adjacent to *FBN2* and *AIM2* confirmed their distinct IFN-response (**Fig. 3D,E**). As expected, the motifs for GATA3 and STAT1 are enriched within constitutively-active and IFN-stimulated MER41-enhancers, respectively (**Fig. 3F**). Accordingly, GATA2/3, MSX2 and co-factors bind the constitutively-active group with higher frequency and intensity (**Fig. 3G,H)**, and in contrast, immune TFs including STAT1 and IRF1 prefer to bind IFN-stimulated enhancers based on the data for multiple human cell types (**Supplemental Fig. S14**) - partly explains why one group of MER41-enhancers are IFN-stimulated while the other group is constitutively-activate in human TSCs. Together, substantial differences regarding the subfamilies and TF binding exist between the placenta- and immune-related MER41-enhancers, suggesting the functional divergence of MER41 elements during evolution.

**Figure 3.**
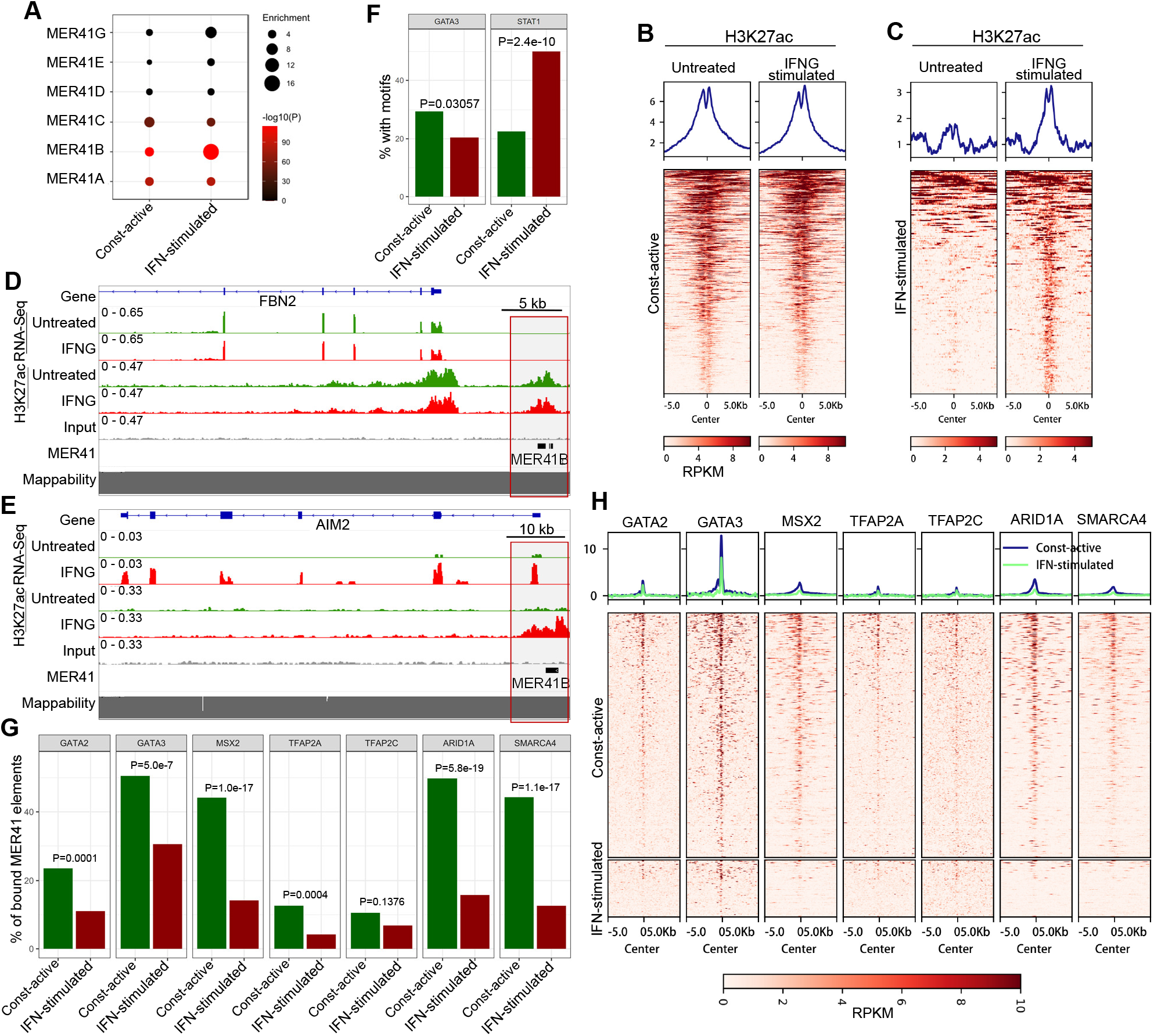
Characterization and comparison of the MER41-enhancers that are constitutively-active or IFN-stimulated in human TSCs. **(A)** Enrichment of MER41 subfamilies in the two groups of MER41-enhancers that are constitutively-active or IFN-stimulated in human TSCs. (**B,C**) Heatmaps show the H3K27ac intensity flanking the two groups of MER41-enhancers with or without IFNG-stimulation. (**D**) Frequency of the motifs for GATA3 and STAT1 in constitutively-active or IFN-stimulated MER41-enhancers, respectively. P-values calculated by using Fisher’s Exact Test are indicated. **(E,F)** Representative IGV tracks show the alterations of gene expression and H3K27ac occupancy flanking the MER41-enhancers adjacent to *FBN2* and *AIM2*, respectively. The color gradients indicate the RPKM values (ChIP – Input) calculated from the ChIP-Seq data. **(G)** Bar plots for the binding frequency of GATA2/3, MSX2 and co-factors on the two groups of MER41-enhancers. P-values calculated by using Fisher’s Exact Test are indicated. **(H)** Heatmaps show the binding intensity of GATA2/3, MSX2 and co-factors on the two groups of MER41-enhancers. The color gradients indicate the RPKM values (ChIP – Input) calculated from the ChIP-Seq data.

### Prevalent binding of GATA2/3 and MSX2 on different families of ERV-enhancers in human TSCs

Given the enriched binding of the aforementioned trophoblast TFs on MER41-enhancers, we are eager to check if other families of ERV-associated enhancers are also bound by these TFs in human TSCs. Globally, GATA2/3, MSX2 and co-factors preferentially occupy ERVs; specifically, 30.0% and 23.8% of the peaks for GATA2 and GATA3 overlap ERV elements, which is remarkably higher than the 8.8% as expected in randomly shuffled peaks across genome (**Fig. 4A**). We further examined their binding on each ERV family, and as expected, MER41A/B/C are among the top enriched (**Fig. 4B**). Interestingly, dozens of other ERV families, such as LTR8, MER11A, MER4B, LTR5_Hs and MER4A1/D1/D/E, are also significantly enriched (**Fig. 4B, Supplemental Fig. S15**). Notably, many of the enriched ERV families are primate- or even human-specific (**Table S5**). Importantly, the ERV families enriched in TSC enhancers and in the peaks for GATA2/3 and MSX2 correlate well (**Fig. 4C**), and correspondingly, 49.2% (7,637 out of 15,504) of the TSC enhancers that overlap ERVs are bound by GATA2/3 or MSX2 (**Fig. 4D**) – significantly higher than the percentage for other enhancers (P<2.2e-16, OddRatio=3.1). Notably, the degree of enriched binding on ERV-derived enhancers is higher for GATA2/3+MSX2 and GATA-only peaks relative to MSX2-only peaks (**Supplemental Fig. S16, S17, S18**). Indeed, the top enriched ERV families not only have global binding by GATA3 and MSX2, but also possess high intensity of H3K27ac marks (**Fig. 4E**) – suggesting these ERV elements also form enhancers. Closer inspection further confirmed the binding of these TFs on one LTR8-associated enhancer adjacent to the imprinted gene *H19* (**Fig. 4F**) and one MER11A-associated enhancer adjacent to *FSTL3* (**Fig. 4G**) – these two genes are both tightly associated with placenta development and diseases (Fowden et al. 2006; Xie et al. 2018; Gong et al. 2021). Together, these results suggest that apart from MER41 elements, these core trophoblast TFs also have prevalent binding on many other families of ERV-associated enhancers – suggesting the broad involvement of these TFs and the bound ERV-associated enhancers in human TSCs.

**Figure 4.**
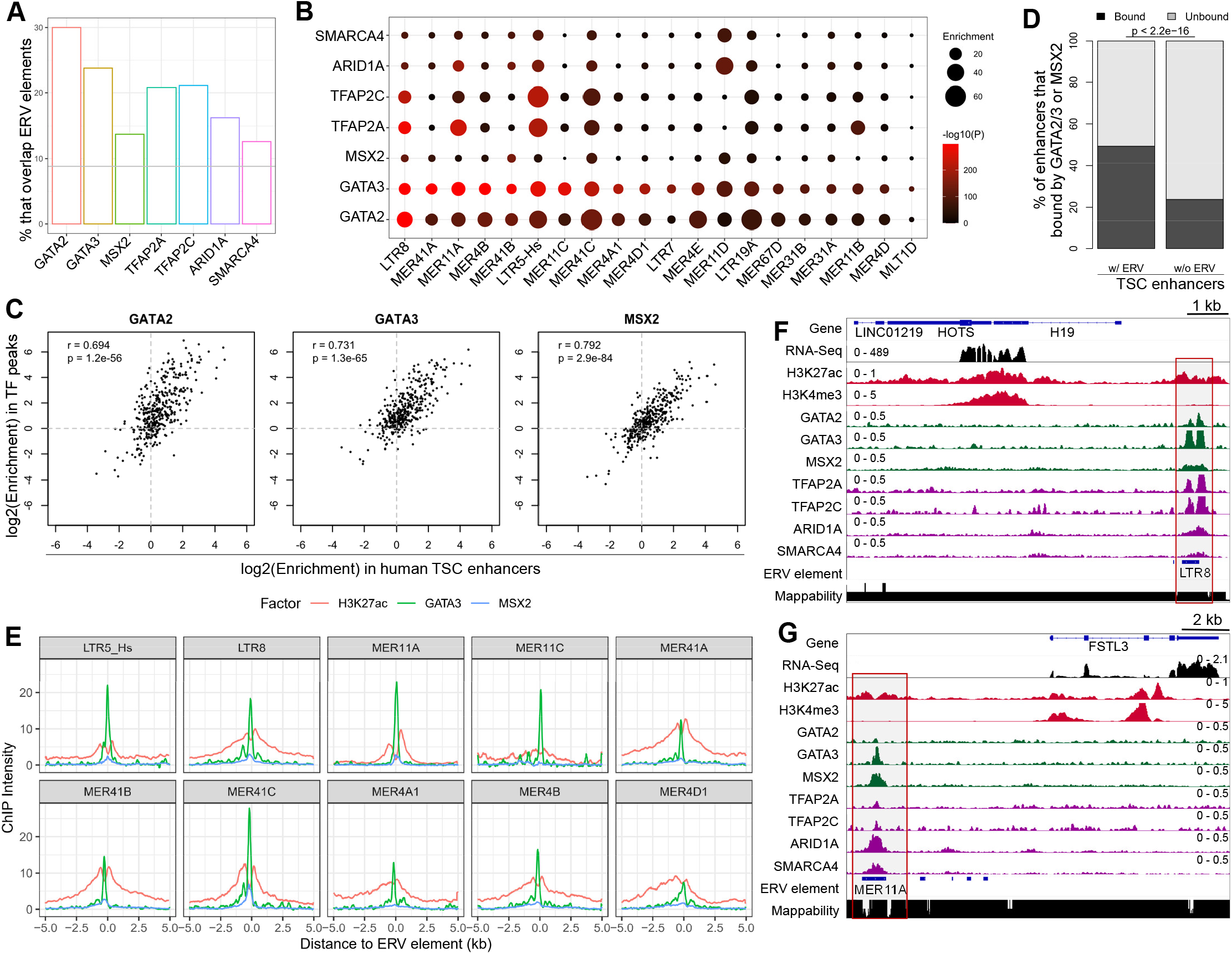
Binding of GATA2/3 and co-factors on different families of ERV-associated enhancers in human TSCs. **(A)** Percentages of the peaks that binding ERV elements for GATA2/3, MSX2 and co-factors. The gray line represents the expected percentage calculated through the comparison of the randomly shuffled peaks (after pooling the peaks for GATA2/3, MSX2 and co-factors) against ERV elements. (**B**) Enrichment of the top twenty ERV families that have enriched binding by GATA2/3, MSX2 or co-factors. These ERV families are selected based on the ranking according to the enrichment p-values against the TF peaks. (**C**) Correlation for the ERV enrichment in annotated enhancers vs. GATA2/3 and MSX2 peaks. **(D)** Comparison of the percentages of enhancers (w/wo ERV overlapping) that are bound by at least one TFs among GATA2/3 and MSX2. P-value calculated by Fisher’s Exact Test is denoted. **(E)** Averaged profiles show the intensity of H3K27ac, GATA3 and MSX2 occupancy on the top ten ERV families that have enriched binding by GATA2/3, MSX2 and co-factors. **(F,G)** Representative IGV tracks show the epigenetic profiles and trophoblast TF binding near the LTR8- and MER11A-associated enhancers adjacent to *H19* and *FSTL3*, respectively.

### MSX2 represses numerous STB-genes in human TSCs through the binding on ERV-associated regulatory elements

Despite the prevalent binding of GATA2/3 and MSX2 on ERV-associated enhancers, it should be noted that these TFs have distinct functions: while GATA2/3 are recognized as pioneer TFs in human TSCs for transcriptional activation (Krendl et al. 2017), MSX2 was recently identified as a transcriptional repressor to restrain the undesired expression of STB-genes in human TSCs (Hornbachner et al. 2021). Given that how ERV elements mediate the function of transcription repressors is rarely reported, in-depth analysis on the binding and function of MSX2 on ERV-associated enhancers is highly desirable. After confirming the repression of STB-genes by MSX2 (**Supplemental Fig. S19**) through re-analysis of public transcriptome data (Hornbachner et al. 2021), we further found that MSX2 has enriched binding on dozens of ERV families, including MER41A/B/C, LTR8, LTR8B, MER21A, and LTR3A (**Fig. 5A**). To clarify the function of MSX2 on ERVs, we compared the H3K27ac marks on the top ten enriched ERV families after the knockdown of MSX2. As expected, knockdown of MSX2 causes increased H3K27ac levels on the MSX2-bound ERV elements, yet the degrees of changes differ among ERV families (**Fig. 5B**). For example, the H3K27ac level for LTR8B is increased more than three folds, while LTR3A is increased more moderately. As control, the H3K27ac level on unbound ERV families shows no remarkable change (**Supplemental Fig. S20**). These results suggest that MSX2 has a global repressive effect on the bound ERV-enhancers in human TSCs.

**Figure 5.**
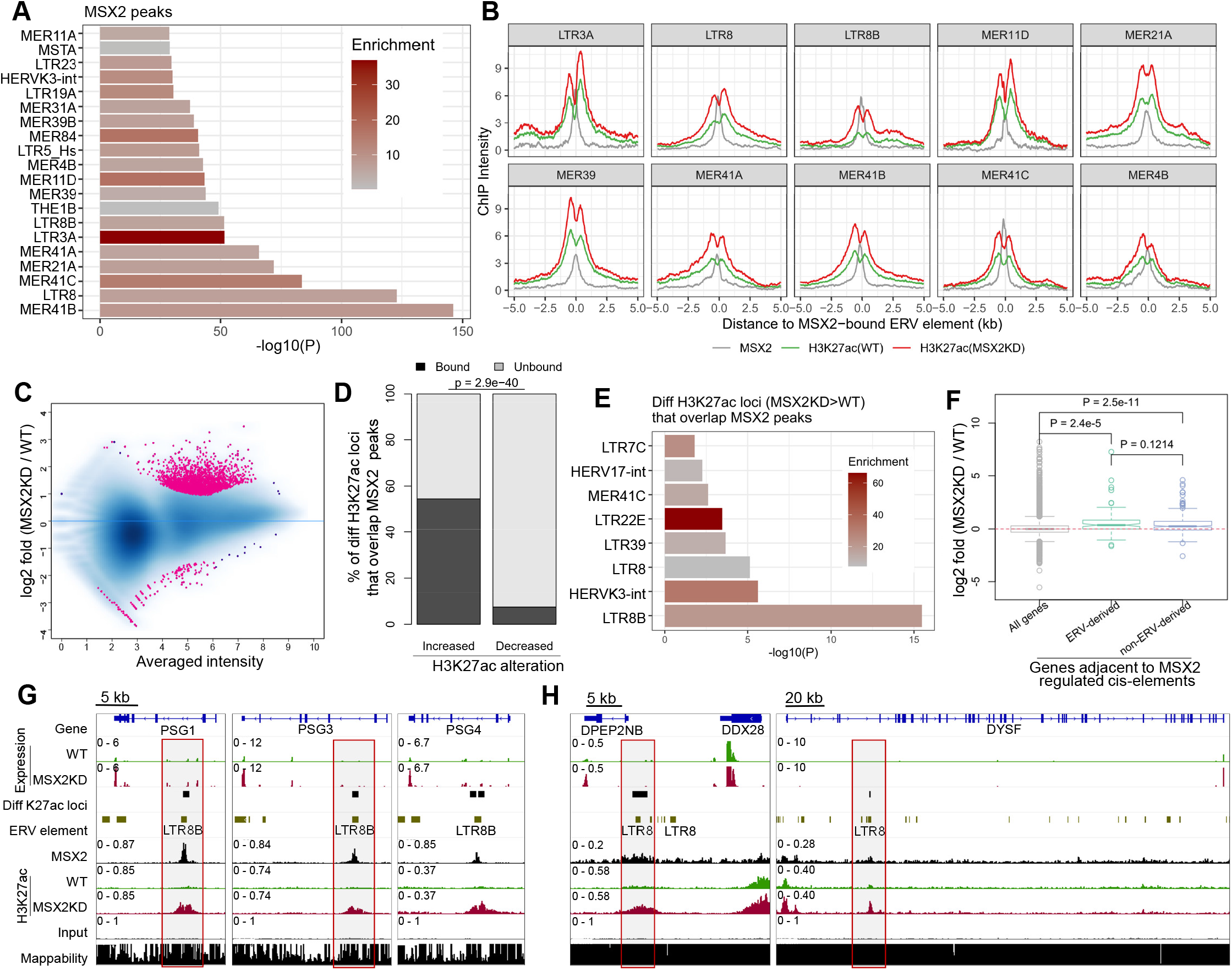
MSX2 restricts the expression of some STB-genes in human TSCs through the binding of ERV-derived enhancers. **(A)** Enrichment of ERV families within MSX2 peaks. The top 20 families as ranked by p-values are presented. **(B)** Averaged curves show the ChIP intensity of MSX2 and H3K27ac (WT vs. MSX2KD) flanking the MSX2-bound elements from the top ten ERV families. **(C)** MA-plot shows the differential H3K27ac peaks between WT and MSX2KD human TSCs. **(D)** Comparison of the MSX2-bound percentages between genomic loci with increased or decreased H3K27ac levels after MSX2KD. P-value calculated with Fisher’s Exact Test is denoted. **(E)** Enrichment of ERV families within MSX2-bound loci that have increased H3K27ac levels after MSX2KD. **(F)** Comparison of the altered expression (MSX2KD vs. WT) across different groups of genes defined based on their association with MSX2-regulated enhancers. A threshold of less than 10 kb from TSSs was used to group genes. MSX2-regulated enhancers are further classified as two groups (ERV-derived or non-ERV-derived) based on their overlapping with ERV elements. P-values calculated from Two-sided Student’s t-test are denoted. **(G,H)** Representative IGV tracks showing several LTR8B- or LTR8-derived enhancers that are likely mediating the MSX2-dependent repression of STB-genes, including *PSG1/3/4, DPEP2NB, DDX28* and *DYSF*.

MSX2 is known to restrict the expression of STB-genes in human TSCs (Hornbachner et al. 2021). To uncover if ERV-associated cis-elements participate in this process, we determined the loci with significantly altered H3K27ac levels after knockdown of MSX2 (MSX2KD) for further analysis by integrating recently published data (Hornbachner et al. 2021). The majority (2,519 out of 2,706) of these differential loci have increased H3K27ac levels (**Fig. 5C**), and MSX2 peaks overlap 54.4% of those with increased H3K27ac – significantly higher than those with decreased H3K27ac (**Fig. 5D**). Several ERV families are also significantly overrepresented within the MSX2-regulated cis-elements (defined as genomic loci that overlap MSX2 peaks and have increased H3K27ac levels in MSX2KD), including LTR8B, HERVK3-int and LTR8 (**Fig. 5E**). We further examined the genes adjacent to MSX2-regulated cis-elements which are further classified as ERV-derived or non-ERV-derived, and as expected, they both have increased expression after MSX2KD (**Fig. 5F**). Manual inspection of the canonical STB-genes repressed by MSX2 indicates that some are likely to be regulated through ERV-derived cis-elements. For example, multiple *PSG* genes (PSG1/3/4/6/8/9) that get derepressed in MSX2KD harbor LTR8B-derived cis-elements that underlie the regulation of MSX2 (**Fig. 5G; Supplemental Fig. S21**). In addition, one LTR8-derived cis-elements between *DPEP2NB* and *DDX28*, and another within *DYSF* are also likely regulated by MSX2 (**Fig. 5H**). Together, these results suggest that MSX2 restricts the undesired expression of many STB-genes in human TSCs through ERV-associated regulatory elements.

## Discussion

In eutherian mammals, the placenta is a transient organ characterized with extraordinary phenotypic diversity (Gerri et al. 2020; Hemberger et al. 2020). At the molecular level, hundreds of placental genes have lineage-specific expression (Dunn-Fletcher et al. 2018; Soncin et al. 2018; Rosenkrantz et al. 2021; Sun et al. 2021), and many are driven by the fast-evolving enhancers which frequently overlap specific ERV families (Chuong et al. 2013; Sun et al. 2021). It is highly desirable to delineate how ERV-associated enhancers participate in the trophoblast gene network, which is under the cooperative regulation of many TFs (Papuchova and Latos 2022).

Our study was initiated by focusing on the primate-specific MER41 family, which is among the top enriched in human placental enhancers (Sun et al. 2021). MER41 elements are also known to create thousands of IFN-stimulated enhancers to drive innate immunity evolution (Schmid and Bucher 2010; Chuong et al. 2016; Raviram et al. 2018). Yet unlike immune-related MER41 elements which have already been well characterized, how MER41-enhancers are regulated in human placenta remains obscure. Interestingly, motif scanning of the consensus and representative sequences of MER41 indicates that they harbors the binding motifs for multiple TFs, including GATA2/3, MSX2 and GRHL2 which are important regulators for trophoblast lineage maintenance and development in human and/or mice (Ralston et al. 2010; Walentin et al. 2015; Home et al. 2017; Krendl et al. 2017; Rhee et al. 2017; Hornbachner et al. 2021). Notably, integration of public data (Krendl et al. 2017; Hornbachner et al. 2021) verified that the majority of MER41-enhancers are bound by GATA3, MSX2 and their co-factors (TFAP2A/2C, ARID1A and SMARCA4). Therefore, the trophoblast TFs, particularly GATA2/3 and MSX2, were identified as promising regulators of MER41-enhancers in human TSCs. Functionally, it is tempting to speculate that, through the binding of GATA2/3, MSX2 and their co-factors, MER41-enhancers particulate in the human trophoblast regulatory network.

While MER41 is among the top enriched in human placental enhancers, dozens of other ERV families are also highly enriched – therefore we were wondering if they are regulated through the same set of TFs. Strikingly, we found that GATA2/3 and MSX2 have prevalent binding on dozens of other families of ERV-associated enhancers, many are primate-specific (eg. LTR8, MER4 and MER11) or even human-specific (eg. LTR5_Hs). Indeed, the ERV families frequently bound by these TFs are well correlated with those enriched in TSC enhancers, suggesting GATA2/3 and MSX2 have broad effect on ERV-associated TSC enhancers. Notably, GATA2/3 are generally recognized as transcriptional activators (Takaku et al. 2016; Krendl et al. 2017), while MSX2 as a transcriptional repressor for STB genes (Hornbachner et al. 2021). In addition, GATA2 and GATA3 are known to have functional redundancy in mouse placenta (Home et al. 2017). Therefore, the cooperation or even competition of these TFs in regulating ERV-associated enhancers is expected. Currently, the molecular functions of these TFs have still not been systematically compared. We expect that the combinatorial function of these TFs on the co-binding loci – including ERV-derived ones – will be clarified through the knockout of each TF and the manipulation of their binding motifs in representative cis-elements.

Even though GATA2/3 and MSX2 are most abundant in placenta, they also express and function in a few other tissues or cell types. For example, GATA3 is also expressed in T cells (Wei et al. 2011; Van de Walle et al. 2016) and mammary gland (Theodorou et al. 2013). So, why the ERV-associated enhancers – which are frequently bound by these TFs – are preferably active in placenta? One possible reason is the unique epigenetic and chromatin signatures owned by placenta. Unlike most other tissues which have global hyper-methylation, placenta is abundant with partially methylated domains (Schroeder et al. 2013; Decato et al. 2017) which are also featured by many tumors (Hansen et al. 2011). Mechanistically, one previous study speculated that placenta and tumor share converged DNA methylation pathways that mediate the establishment of such epigenetic features (Lorincz and Schubeler 2017). Given the well-recognized function of DNA methylation for epigenetic repression, it is possible that many genomic loci (particularly ERV-derived ones) are more accessible in the placenta relative to other tissues – thus these loci are more likely to be activated in placenta. The derepression of ERVs has also been observed in numerous types of tumors, presumably due to the attenuation of the epigenetic silencing (Chuong et al. 2017; Ito et al. 2020). Therefore, it is possible that the prevalent binding of trophoblast TFs on ERV-associated enhancers may also rely on the relatively relaxed epigenetic environment in trophoblast cells. Apart from the epigenetic/chromatin environment, additional mechanisms, such as the existence of specific co-factors, may also contribute to the tissue-specific binding and regulation of ERV-associated enhancers by these TFs in placenta.

Most previous studies about ERV-derived enhancers focused on their activation, yet how they mediate the function of transcription repressors is rarely reported. Aiming at mechanistic insights into the involvement of ERVs for transcriptional repression, we performed in-depth analysis on MSX2, a homeobox gene which is known to be crucial in restraining the undesired expression of STB-genes in human TSCs (Hornbachner et al. 2021). Notably, while hundreds of KRAB zinc finger proteins are known to be evolved to repress ERVs in mammals (Imbeault et al. 2017; Wolf et al. 2020), how homeobox genes participate in the repression of ERVs remains poorly understood. Interestingly, the knockdown of MSX2 resulted in globally increased H3K27ac levels on the ERV-associated enhancers it bound, which suggests that it has global repressive effect on the bound ERVs. Importantly, for the genomic loci under the repression of MSX2, several ERV families are also significantly overrepresented. Genes adjacent to MSX2-bound enhancers – no matter they are ERV-derived or not - usually have increased expression after MSX2KD. Furthermore, some STB-genes are likely to be regulated through ERV-derived enhancers, such as LTR8B-derived enhancers for multiple *PSG* genes, an LTR8-derived adjacent to *DPEP2NB, DDX28* and *DYSF*. Therefore, MSX2 restricts the undesired expression of many STB-genes in human TSCs through ERV-associated enhancers. These results further suggest that through the recruiting of different transcriptional activators and repressors, ERVs can be co-opted as a cis-regulatory module to mediate the accurate expression of associated genes in the host.

Taken together, this study uncovered the prevalent regulation of multiple trophoblast TFs, particularly GATA2/3, MSX2 and their co-factors, on numerous ERV-derived enhancers in human TSCs. Through comprehensive analysis on the links between ERV-derived enhancers, upstream regulators and downstream targets, this study provides novel mechanistic insights into the functional involvement of ERVs in the human trophoblast regulatory network.

## Materials and methods

### Human TSC culture and treatment

Human TSCs derived from human cytotrophoblast cells (Okae et al. 2018) were a gift from the Okae lab. They were cultured in trophoblast stem cell medium (TSM), as described previously (Sun et al. 2021). Interferon stimulation was performed by adding 1000U/mL of IFNG (PBL, 11500-2) into the culturing medium of human TSCs. After 24 hours, untreated and IFNG treated samples were both collected for subsequent ChIP-Seq (H3K27ac) and RNA-Seq experiments.

### RNA-Seq

Total RNA for human TSCs was extracted using RNeasy Micro kit (Qiagen, 74004) with on-column DNase digestion, and then submitted for library construction by TruSeq stranded mRNA sample preparation kit (Illumina). Two biological replicates were used per sample. RNA-Seq libraries were sequenced as 75 bp paired-end reads with HiSeq2500 (Illumina) platform.

Raw reads were trimmed with Trim Galore v0.6.4 (https://github.com/FelixKrueger/TrimGalore). Transcript Per Million (TPM) values were calculated with RSEM v1.3.2 (Li and Dewey 2011). To perform differential expression analysis, we aligned trimmed reads to the reference genome (GRCh38 for human) using STAR v2.7.3 (Dobin et al. 2013), and then obtained gene-level read counts using the *featureCount* function from subread v2.0.0 (Liao et al. 2013). At last, differentially expressed genes were identified using DESeq2 v1.30.1 (Love et al. 2014) with the cutoff: FDR<0.05 and |log2Foldchange|>1.

### ChIP-Seq

ChIP-Seq for human TSCs was performed following our previous study (Sun et al. 2021). Chromatin fragmentation was performed using Diagenode Bioruptor Plus Sonicator. The antibody for H3K27ac (Abcam, ab4729) is used. The amount of chromatin is 20 μg per reaction. Two biological replicates were used per sample. ChIP-Seq libraries were constructed using Takara SMARTer ThruPLEX DNA-Seq Kit (Takara, R400674), and sequenced as 50 bp paired-end reads with HiSeq2500 (Illumina) platform.

Reads were trimmed with TrimGalore v0.6.4 and then aligned to the corresponding reference genome (GRCh38 for human) using Bowtie v2.3.5 (Langmead and Salzberg 2012) with default settings. PCR duplicates were removed using the *rmdup* function of samtools v1.13 (Li et al. 2009). After confirming the data reproducibility, reads from biological replicates were pooled together for further analysis. Peak calling was performed with MACS v2.2.6 (Zhang et al. 2008). The peaks were further cleaned by removing those that overlap ENCODE Blacklist V2 regions (Amemiya et al. 2019). Differential binding analysis was performed using DiffBind v3.4.11 (Ross-Innes et al. 2012) with settings: minOverlap = 1, summits = 400, method = DBA_EDGER.

### Reference genome and annotation

Reference genome and gene annotation for human (GRCh38) were downloaded from the ENSEMBL database (release 102) (Yates et al. 2020). Transposable element annotations were downloaded from the RepeatMasker website (http://www.repeatmasker.org/) on May 27, 2016. The clades for ERV families were obtained from the Dfam database (Hubley et al. 2016). Genome mappability along the reference genome was calculated using the GEM-mappability program from GEM (GEnome Multitool) suite (Derrien et al. 2012).

### Gene Ontology enrichment analysis

Gene Ontology enrichment analyses for differentially expressed genes were performed using Metascape (Zhou et al. 2019). Gene Ontology enrichment analyses for genomic regions (eg. peaks and putative enhancers) were performed with GREAT (McLean et al. 2010).

### Motif analysis

Motif scanning on MER41 consensus sequences (downloaded from the RepeatMasker website on May 27, 2016) were performed against the non-redundant JASPAR2022 CORE vertebrates motif database (JASPAR2022_CORE_vertebrates_non-redundant) using FIMO (Grant et al. 2011) with settings: --text --thresh 1e-3. The locations of predicted motifs were visualized with IGV v2.11.1 (Thorvaldsdottir et al. 2013), and redundant predictions (eg. multiple prediction results of the same TF on overlapped position) were manually adjusted to get optimized visualization. Motif occurrence on representative MER41 elements was retrieved directly from UCSC Genome Browser (Lee et al. 2020). Motif enrichment analysis on MER41-derived enhancers relative to MER41-elements that lacking H3K27ac mark was performed by using the findMotifs.pl script from HOMER v4.11 (Heinz et al. 2010) with default settings.

### Epigenetic-annotation of regulatory elements

Putative regulatory elements are defined based on histone modifications and genomic distribution. Promoters are defined as H3K4me3 occupied regions, and enhancers as H3K27ac peaks that are more than 500 bp away from TSSs.

### Annotation of the genes adjacent to given genomic regions

The genes adjacent to given genomic regions (eg. peaks, enhancers, ERV elements) were determined by comparing gene TSSs against specified genomic regions using the *window* function of BEDtools v2.29.2 (Quinlan and Hall 2010), with the distance threshold specified by the papameter *w*. TSS annotation file was retrieved from ENSEMBL database (release 102) by using BioMart (Yates et al. 2020).

### Differential expression analysis between human and mouse

Interspecies differential expression analysis between human and mouse placentae were performed using ExprX (https://github.com/mingansun/ExprX) as previous described (Sun et al. 2021). In brief, only 1-to-1 orthologous gene pairs annotated by ENSEMBL database (Yates et al. 2020) were used for analysis, and differentially expressed genes were determined using Rank Product method with the cutoff of FDR < 0.05 and |log2foldchange| > 1.

### ERV enrichment analysis

To determine if certain ERV families are overrepresented within given genomic regions (eg. enhancers or peaks), we adopted the *fisher* function of BEDtools v2.29.2 (Quinlan and Hall 2010), which determines the enrichment fold and p-value between two lists of genomic intervals by using Fisher’s Exact Test. A wrapper named TEenrich-FET (https://github.com/mingansun/TE_analysis) was further implemented to simplify the analysis of multiple ERV families. To control for Family-Wise Error Rate, the calculated p-values were further adjusted with Bonferroni method.

### Statistical analysis and data visualization

All statistical analyses were performed with R statistical programming language (Team 2020). Heatmaps for ChIP-Seq data were generated using DeepTools v3.5.1 (Ramirez et al. 2014), with ChIP signal normalized as RPKM values. Heatmaps from gene expression clustering analysis were generated using pheatmap (https://github.com/raivokolde/pheatmap). RNA-Seq and ChIP-Seq tracks were visualized using IGV v2.11.1 (Thorvaldsdottir et al. 2013).

## Data access

All raw and processed sequencing data generated in this study have been submitted to the NCBI Gene Expression Omnibus (GEO; https://www.ncbi.nlm.nih.gov/geo/) under accession number GSE209541.

## Acknowledgements

This study was supported by the grants from the National Natural Science Foundation of China (32270584 & 31900422 to MAS; 82130047 & 81971414 to BC), the National Key Research and Development Program of China (2022YFC2702403 to BC), the Intramural Research Program of the NICHD, NIH (TSM), the 111 Project D18007, the Priority Academic Program Development of Jiangsu Higher Education Institutions (PAPD), and the Natural Sciences Foundation of Fujian Province of China (2020J06003 to BC). We thank Dr. Hiroaki Okae and Dr. Takahiro Arima for kindly providing human trophoblast stem cells. This study utilized the computational resources of Yangzhou University College of Veterinary Medicine High-Performance Computing cluster.

## References

Amemiya HM, Kundaje A, Boyle AP. 2019. The ENCODE Blacklist: Identification of Problematic Regions of the Genome. Sci Rep 9: 9354.

Ander SE, Diamond MS, Coyne CB. 2019. Immune responses at the maternal-fetal interface. Sci Immunol 4.

Buttler CA, Chuong EB. 2022. Emerging roles for endogenous retroviruses in immune epigenetic regulation. Immunol Rev 305: 165–178.

Chuong EB, Elde NC, Feschotte C. 2016. Regulatory evolution of innate immunity through co-option of endogenous retroviruses. Science 351: 1083–1087.

Chuong EB, Elde NC, Feschotte C. 2017. Regulatory activities of transposable elements: from conflicts to benefits. Nat Rev Genet 18: 71–86.

Chuong EB, Rumi MA, Soares MJ, Baker JC. 2013. Endogenous retroviruses function as species-specific enhancer elements in the placenta. Nat Genet 45: 325–329.

Decato BE, Lopez-Tello J, Sferruzzi-Perri AN, Smith AD, Dean MD. 2017. DNA Methylation Divergence and Tissue Specialization in the Developing Mouse Placenta. Mol Biol Evol 34: 1702–1712.

Deniz O, Frost JM, Branco MR. 2019. Regulation of transposable elements by DNA modifications. Nat Rev Genet 20: 417–431.

Derrien T, Estelle J, Marco Sola S, Knowles DG, Raineri E, Guigo R, Ribeca P. 2012. Fast computation and applications of genome mappability. PLoS One 7: e30377.

Dobin A, Davis CA, Schlesinger F, Drenkow J, Zaleski C, Jha S, Batut P, Chaisson M, Gingeras TR. 2013. STAR: ultrafast universal RNA-seq aligner. Bioinformatics 29: 15–21.

Dunn-Fletcher CE, Muglia LM, Pavlicev M, Wolf G, Sun MA, Hu YC, Huffman E, Tumukuntala S, Thiele K, Mukherjee A et al. 2018. Anthropoid primate-specific retroviral element THE1B controls expression of CRH in placenta and alters gestation length. PLoS Biol 16: e2006337.

Fowden AL, Sibley C, Reik W, Constancia M. 2006. Imprinted genes, placental development and fetal growth. Horm Res 65 Suppl 3: 50–58.

Gerri C, McCarthy A, Alanis-Lobato G, Demtschenko A, Bruneau A, Loubersac S, Fogarty NME, Hampshire D, Elder K, Snell P et al. 2020. Initiation of a conserved trophectoderm program in human, cow and mouse embryos. Nature 587: 443–447.

Gong S, Gaccioli F, Dopierala J, Sovio U, Cook E, Volders PJ, Martens L, Kirk PDW, Richardson S, Smith GCS et al. 2021. The RNA landscape of the human placenta in health and disease. Nat Commun 12: 2639.

Grant CE, Bailey TL, Noble WS. 2011. FIMO: scanning for occurrences of a given motif. Bioinformatics 27: 1017–1018.

Hansen KD, Timp W, Bravo HC, Sabunciyan S, Langmead B, McDonald OG, Wen B, Wu H, Liu Y, Diep D et al. 2011. Increased methylation variation in epigenetic domains across cancer types. Nat Genet 43: 768–775.

Heinz S, Benner C, Spann N, Bertolino E, Lin YC, Laslo P, Cheng JX, Murre C, Singh H, Glass CK. 2010. Simple combinations of lineage-determining transcription factors prime cis-regulatory elements required for macrophage and B cell identities. Mol Cell 38: 576–589.

Hemberger M, Hanna CW, Dean W. 2020. Mechanisms of early placental development in mouse and humans. Nat Rev Genet 21: 27–43.

Home P, Kumar RP, Ganguly A, Saha B, Milano-Foster J, Bhattacharya B, Ray S, Gunewardena S, Paul A, Camper SA et al. 2017. Genetic redundancy of GATA factors in the extraembryonic trophoblast lineage ensures the progression of preimplantation and postimplantation mammalian development. Development 144: 876–888.

Hornbachner R, Lackner A, Papuchova H, Haider S, Knofler M, Mechtler K, Latos PA. 2021. MSX2 safeguards syncytiotrophoblast fate of human trophoblast stem cells. Proc Natl Acad Sci U S A 118.

Hubley R, Finn RD, Clements J, Eddy SR, Jones TA, Bao W, Smit AF, Wheeler TJ. 2016. The Dfam database of repetitive DNA families. Nucleic Acids Res 44: D81–89.

Imbeault M, Helleboid PY, Trono D. 2017. KRAB zinc-finger proteins contribute to the evolution of gene regulatory networks. Nature 543: 550–554.

Ito J, Kimura I, Soper A, Coudray A, Koyanagi Y, Nakaoka H, Inoue I, Turelli P, Trono D, Sato K. 2020. Endogenous retroviruses drive KRAB zinc-finger protein family expression for tumor suppression. Sci Adv 6.

Johnson WE. 2019. Origins and evolutionary consequences of ancient endogenous retroviruses. Nat Rev Microbiol 17: 355–370.

Kojima KK. 2018. Human transposable elements in Repbase: genomic footprints from fish to humans. Mob DNA 9: 2.

Krendl C, Shaposhnikov D, Rishko V, Ori C, Ziegenhain C, Sass S, Simon L, Muller NS, Straub T, Brooks KE et al. 2017. GATA2/3-TFAP2A/C transcription factor network couples human pluripotent stem cell differentiation to trophectoderm with repression of pluripotency. Proc Natl Acad Sci U S A 114: E9579–E9588.

Langmead B, Salzberg SL. 2012. Fast gapped-read alignment with Bowtie 2. Nat Methods 9: 357–359.

Lee CM, Barber GP, Casper J, Clawson H, Diekhans M, Gonzalez JN, Hinrichs AS, Lee BT, Nassar LR, Powell CC et al. 2020. UCSC Genome Browser enters 20th year. Nucleic Acids Res 48: D756–D761.

Li B, Dewey CN. 2011. RSEM: accurate transcript quantification from RNA-Seq data with or without a reference genome. BMC Bioinformatics 12: 323.

Li H, Handsaker B, Wysoker A, Fennell T, Ruan J, Homer N, Marth G, Abecasis G, Durbin R, Genome Project Data Processing S. 2009. The Sequence Alignment/Map format and SAMtools. Bioinformatics 25: 2078–2079.

Liao Y, Smyth GK, Shi W. 2013. The Subread aligner: fast, accurate and scalable read mapping by seed-and-vote. Nucleic Acids Res 41: e108.

Lorincz MC, Schubeler D. 2017. Evidence for Converging DNA Methylation Pathways in Placenta and Cancer. Dev Cell 43: 257–258.

Love MI, Huber W, Anders S. 2014. Moderated estimation of fold change and dispersion for RNA-seq data with DESeq2. Genome Biol 15: 550.

Macfarlan TS, Gifford WD, Driscoll S, Lettieri K, Rowe HM, Bonanomi D, Firth A, Singer O, Trono D, Pfaff SL. 2012. Embryonic stem cell potency fluctuates with endogenous retrovirus activity. Nature 487: 57–63.

Maltepe E, Fisher SJ. 2015. Placenta: the forgotten organ. Annu Rev Cell Dev Biol 31: 523–552.

McLean CY, Bristor D, Hiller M, Clarke SL, Schaar BT, Lowe CB, Wenger AM, Bejerano G. 2010. GREAT improves functional interpretation of cis-regulatory regions. Nat Biotechnol 28: 495–501.

Mossman HW. 1987. Vertebrate fetal membranes : comparative ontogeny and morphology, evolution, phylogenetic significance, basic functions, research opportunities. New Brunswick, N.J. : Rutgers University Press.

Okae H, Toh H, Sato T, Hiura H, Takahashi S, Shirane K, Kabayama Y, Suyama M, Sasaki H, Arima T. 2018. Derivation of Human Trophoblast Stem Cells. Cell Stem Cell 22: 50–63 e56.

Papuchova H, Latos PA. 2022. Transcription factor networks in trophoblast development. Cell Mol Life Sci 79: 337.

Paul S, Home P, Bhattacharya B, Ray S. 2017. GATA factors: Master regulators of gene expression in trophoblast progenitors. Placenta 60 Suppl 1: S61–S66.

Quinlan AR, Hall IM. 2010. BEDTools: a flexible suite of utilities for comparing genomic features. Bioinformatics 26: 841–842.

Ralston A, Cox BJ, Nishioka N, Sasaki H, Chea E, Rugg-Gunn P, Guo G, Robson P, Draper JS, Rossant J. 2010. Gata3 regulates trophoblast development downstream of Tead4 and in parallel to Cdx2. Development 137: 395–403.

Ramirez F, Dundar F, Diehl S, Gruning BA, Manke T. 2014. deepTools: a flexible platform for exploring deep-sequencing data. Nucleic Acids Res 42: W187–191.

Ramsey EM, Houston ML, Harris JW. 1976. Interactions of the trophoblast and maternal tissues in three closely related primate species. Am J Obstet Gynecol 124: 647–652.

Raviram R, Rocha PP, Luo VM, Swanzey E, Miraldi ER, Chuong EB, Feschotte C, Bonneau R, Skok JA. 2018. Analysis of 3D genomic interactions identifies candidate host genes that transposable elements potentially regulate. Genome Biol 19: 216.

Rhee C, Lee BK, Beck S, LeBlanc L, Tucker HO, Kim J. 2017. Mechanisms of transcription factor-mediated direct reprogramming of mouse embryonic stem cells to trophoblast stem-like cells. Nucleic Acids Res 45: 10103–10114.

Rosenkrantz JL, Gaffney JE, Roberts VHJ, Carbone L, Chavez SL. 2021. Transcriptomic analysis of primate placentas and novel rhesus trophoblast cell lines informs investigations of human placentation. BMC Biol 19: 127.

Ross-Innes CS, Stark R, Teschendorff AE, Holmes KA, Ali HR, Dunning MJ, Brown GD, Gojis O, Ellis IO, Green AR et al. 2012. Differential oestrogen receptor binding is associated with clinical outcome in breast cancer. Nature 481: 389–393.

Schmid CD, Bucher P. 2010. MER41 repeat sequences contain inducible STAT1 binding sites. PLoS One 5: e11425.

Schoenfelder S, Mifsud B, Senner CE, Todd CD, Chrysanthou S, Darbo E, Hemberger M, Branco MR. 2018. Divergent wiring of repressive and active chromatin interactions between mouse embryonic and trophoblast lineages. Nat Commun 9: 4189.

Schroeder DI, Blair JD, Lott P, Yu HO, Hong D, Crary F, Ashwood P, Walker C, Korf I, Robinson WP et al. 2013. The human placenta methylome. Proc Natl Acad Sci U S A 110: 6037–6042.

Senft AD, Macfarlan TS. 2021. Transposable elements shape the evolution of mammalian development. Nat Rev Genet 22: 691–711.

Sharif J, Endo TA, Nakayama M, Karimi MM, Shimada M, Katsuyama K, Goyal P, Brind’Amour J, Sun MA, Sun Z et al. 2016. Activation of Endogenous Retroviruses in Dnmt1(-/-) ESCs Involves Disruption of SETDB1-Mediated Repression by NP95 Binding to Hemimethylated DNA. Cell Stem Cell 19: 81–94.

Soncin F, Khater M, To C, Pizzo D, Farah O, Wakeland A, Arul Nambi Rajan K, Nelson KK, Chang CW, Moretto-Zita M et al. 2018. Comparative analysis of mouse and human placentae across gestation reveals species-specific regulators of placental development. Development 145.

Stoye JP. 2012. Studies of endogenous retroviruses reveal a continuing evolutionary saga. Nat Rev Microbiol 10: 395–406.

Sun MA, Wolf G, Wang Y, Senft AD, Ralls S, Jin J, Dunn-Fletcher CE, Muglia LJ, Macfarlan TS. 2021. Endogenous Retroviruses Drive Lineage-Specific Regulatory Evolution across Primate and Rodent Placentae. Mol Biol Evol 38: 4992–5004.

Takaku M, Grimm SA, Shimbo T, Perera L, Menafra R, Stunnenberg HG, Archer TK, Machida S, Kurumizaka H, Wade PA. 2016. GATA3-dependent cellular reprogramming requires activation-domain dependent recruitment of a chromatin remodeler. Genome Biol 17: 36.

Team RC. 2020. R: A language and environment for statistical computing. R Foundation for Statistical Computing, Vienna, Austria.

Theodorou V, Stark R, Menon S, Carroll JS. 2013. GATA3 acts upstream of FOXA1 in mediating ESR1 binding by shaping enhancer accessibility. Genome Res 23: 12–22.

Thorvaldsdottir H, Robinson JT, Mesirov JP. 2013. Integrative Genomics Viewer (IGV): high-performance genomics data visualization and exploration. Brief Bioinform 14: 178–192.

Todd CD, Deniz O, Taylor D, Branco MR. 2019. Functional evaluation of transposable elements as enhancers in mouse embryonic and trophoblast stem cells. Elife 8.

Van de Walle I, Dolens AC, Durinck K, De Mulder K, Van Loocke W, Damle S, Waegemans E, De Medts J, Velghe I, De Smedt M et al. 2016. GATA3 induces human T-cell commitment by restraining Notch activity and repressing NK-cell fate. Nat Commun 7: 11171.

Walentin K, Hinze C, Werth M, Haase N, Varma S, Morell R, Aue A, Potschke E, Warburton D, Qiu A et al. 2015. A Grhl2-dependent gene network controls trophoblast branching morphogenesis. Development 142: 1125–1136.

Wei G, Abraham BJ, Yagi R, Jothi R, Cui K, Sharma S, Narlikar L, Northrup DL, Tang Q, Paul WE et al. 2011. Genome-wide analyses of transcription factor GATA3-mediated gene regulation in distinct T cell types. Immunity 35: 299–311.

Wolf G, de Iaco A, Sun MA, Bruno M, Tinkham M, Hoang D, Mitra A, Ralls S, Trono D, Macfarlan TS. 2020. KRAB-zinc finger protein gene expansion in response to active retrotransposons in the murine lineage. Elife 9.

Xie J, Xu Y, Wan L, Wang P, Wang M, Dong M. 2018. Involvement of follistatin-like 3 in preeclampsia. Biochem Biophys Res Commun 506: 692–697.

Yates AD, Achuthan P, Akanni W, Allen J, Allen J, Alvarez-Jarreta J, Amode MR, Armean IM, Azov AG, Bennett R et al. 2020. Ensembl 2020. Nucleic Acids Res 48: D682–D688.

Yu Y, He JH, Hu LL, Jiang LL, Fang L, Yao GD, Wang SJ, Yang Q, Guo Y, Liu L et al. 2020. Placensin is a glucogenic hormone secreted by human placenta. EMBO Rep 21: e49530.

Zhang Y, Liu T, Meyer CA, Eeckhoute J, Johnson DS, Bernstein BE, Nusbaum C, Myers RM, Brown M, Li W et al. 2008. Model-based analysis of ChIP-Seq (MACS). Genome Biol 9: R137.

Zhou Y, Zhou B, Pache L, Chang M, Khodabakhshi AH, Tanaseichuk O, Benner C, Chanda SK. 2019. Metascape provides a biologist-oriented resource for the analysis of systems-level datasets. Nat Commun 10: 1523.

