## Supplemental Figs S1-S21 for "Prevalent regulation of GATA2/3 and MSX2 on endogenous retrovirus-derived regulatory elements in human trophoblast stem cells"

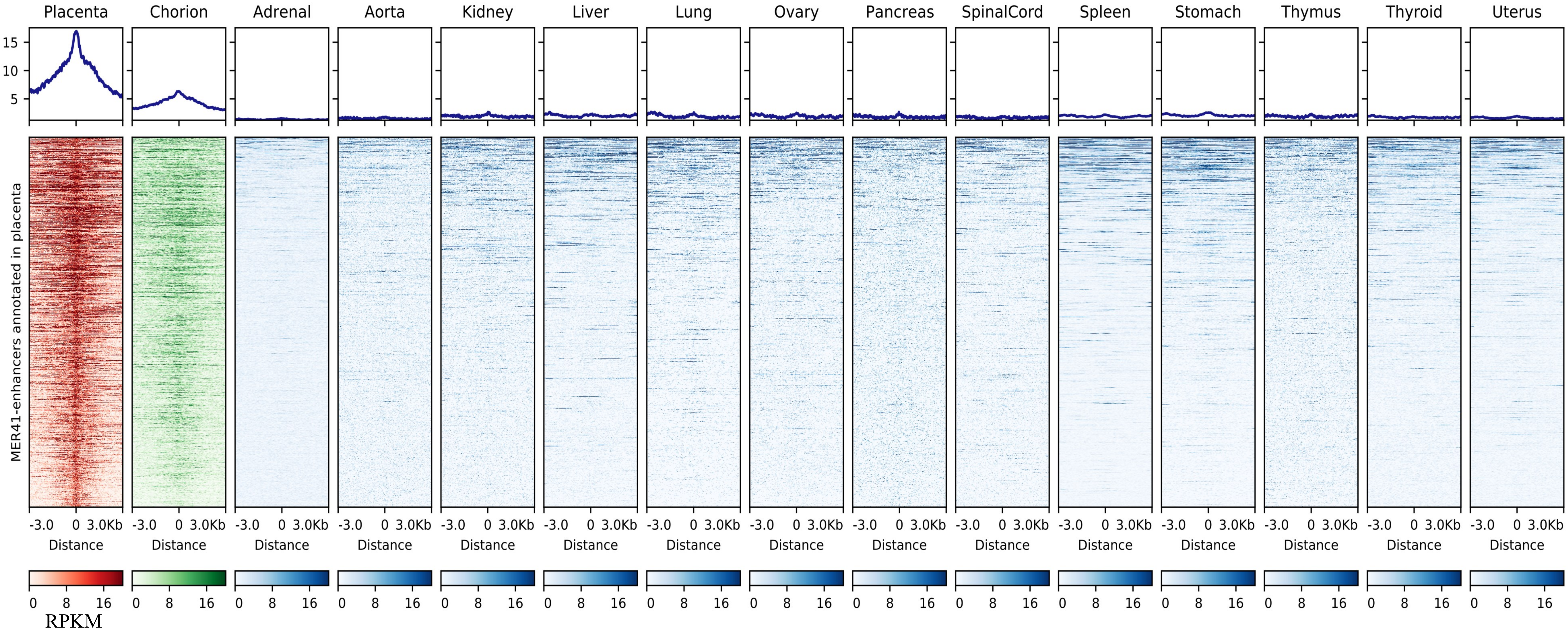


**Fig. S1 Epigenomic profiles in different tissues for the MER41-enhancers annotated in human placenta**
The heatmaps visualize the epigenomic profiles across these tissues for the MER41-enhancers annotated in human placenta. The data confirm that MER41-enhancers are specifically active in placenta (and chorion which is a part of placenta relative to other human tissues. This figure is based on re-analysis of the H3K27ac ChIP-Seq data retrieved from ENCODE project (Table S1). The color gradients indicate the RPKM values calculated from H3K27ac ChIP-Seq data.

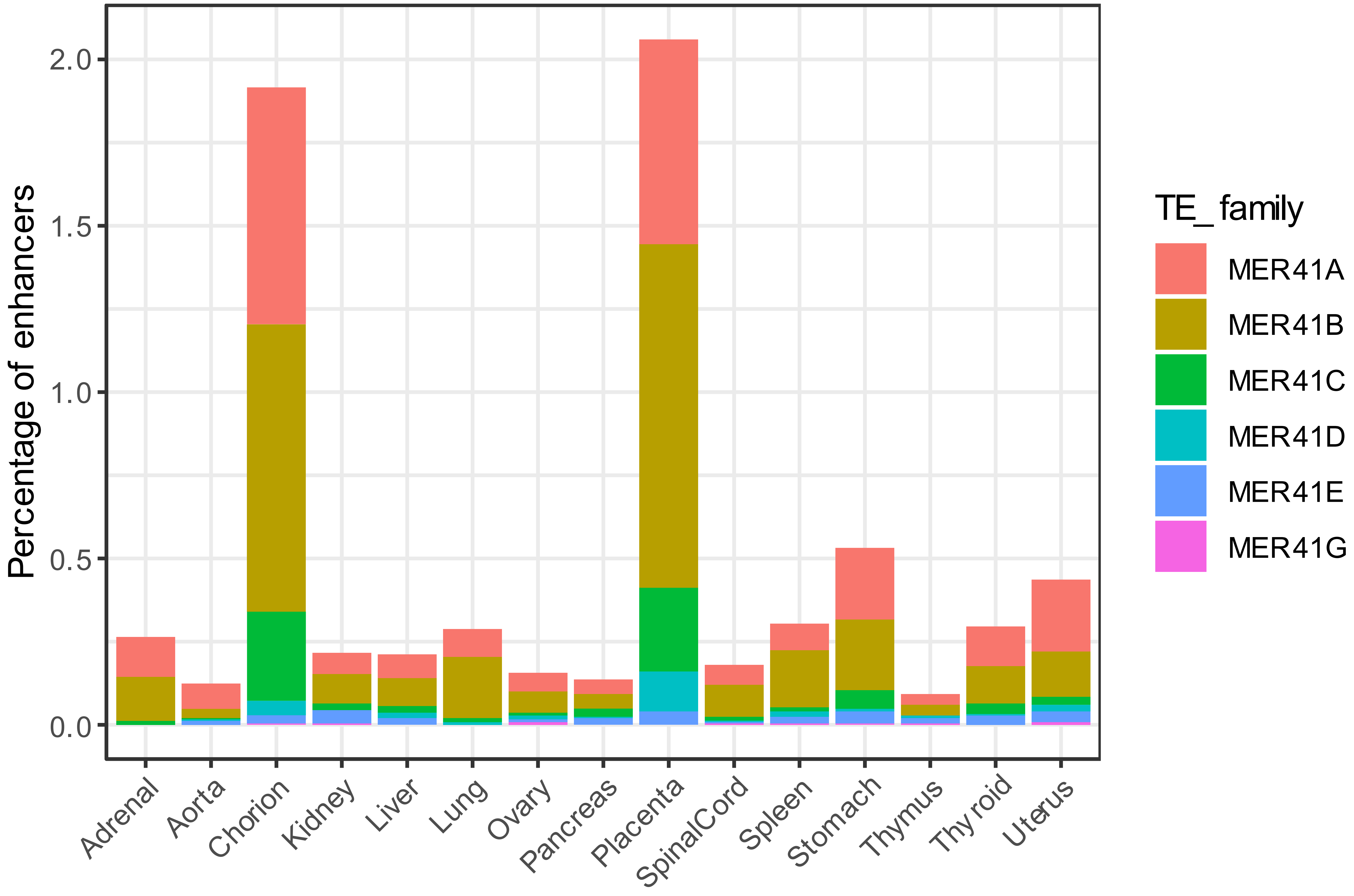


**Fig. S2 Percentages of enhancers for different tissues that overlap MER41 elements**

The bar plots compare the percentages of enhancers for placenta, chorion and other human tissues that overlap the six MER41 subfamilies (A/B/C/D/E/G). The result indicates that placenta- and chorion-enhancers harbor higher percentages of MER41-derived enhancers relative to other tissues.

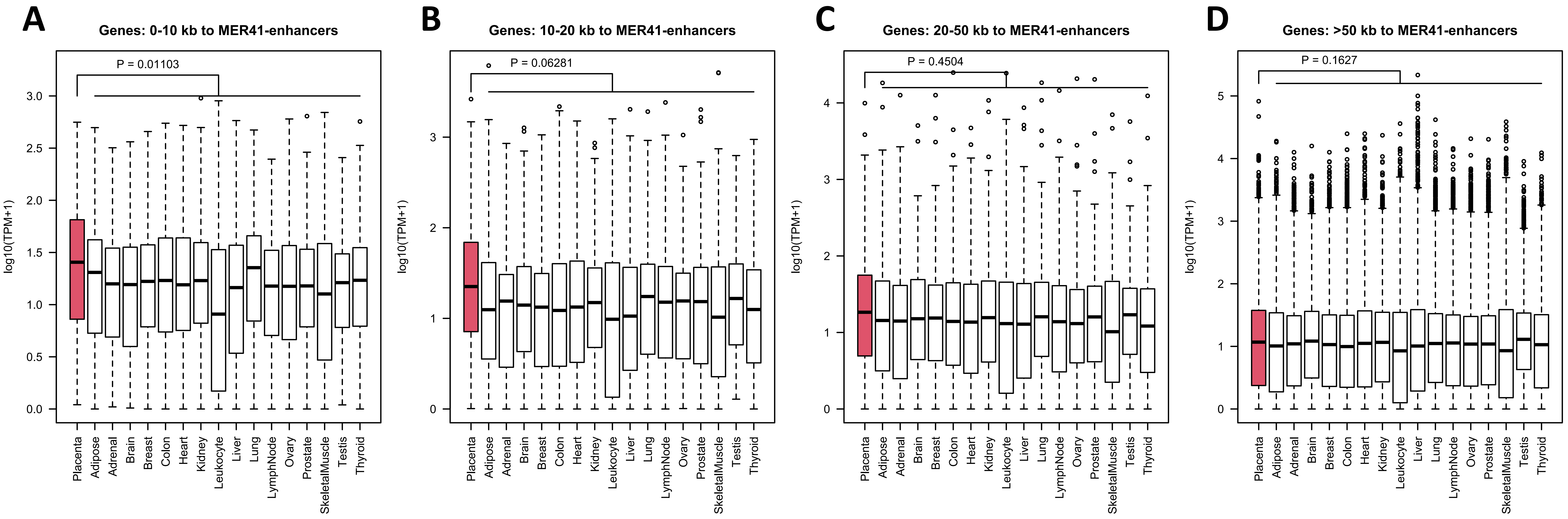


**Fig. S3 Expression profiles for genes adjacent or distal to MER41-enhancers among different human tissues**

The boxplots compare the expression levels of the genes adjacent or distal to placental MER41-enhancer among different human tissues. MER41-enhancer adjacent genes (**A, B, C**) are defined as different groups based on the distance threshold (ie. 0-10kb, 10-20kb, 20-50kb) to TSSs. Genes distal to placental MER41-enhancers (**D**) are annotated as genes with their TSSs more than 50 kb away. This figure is based on re-analysis of transcriptomic data retrieved from BodyMap 2.0 (Table S1). P-values calculated by using one-tailed Student's t-test are indicated.

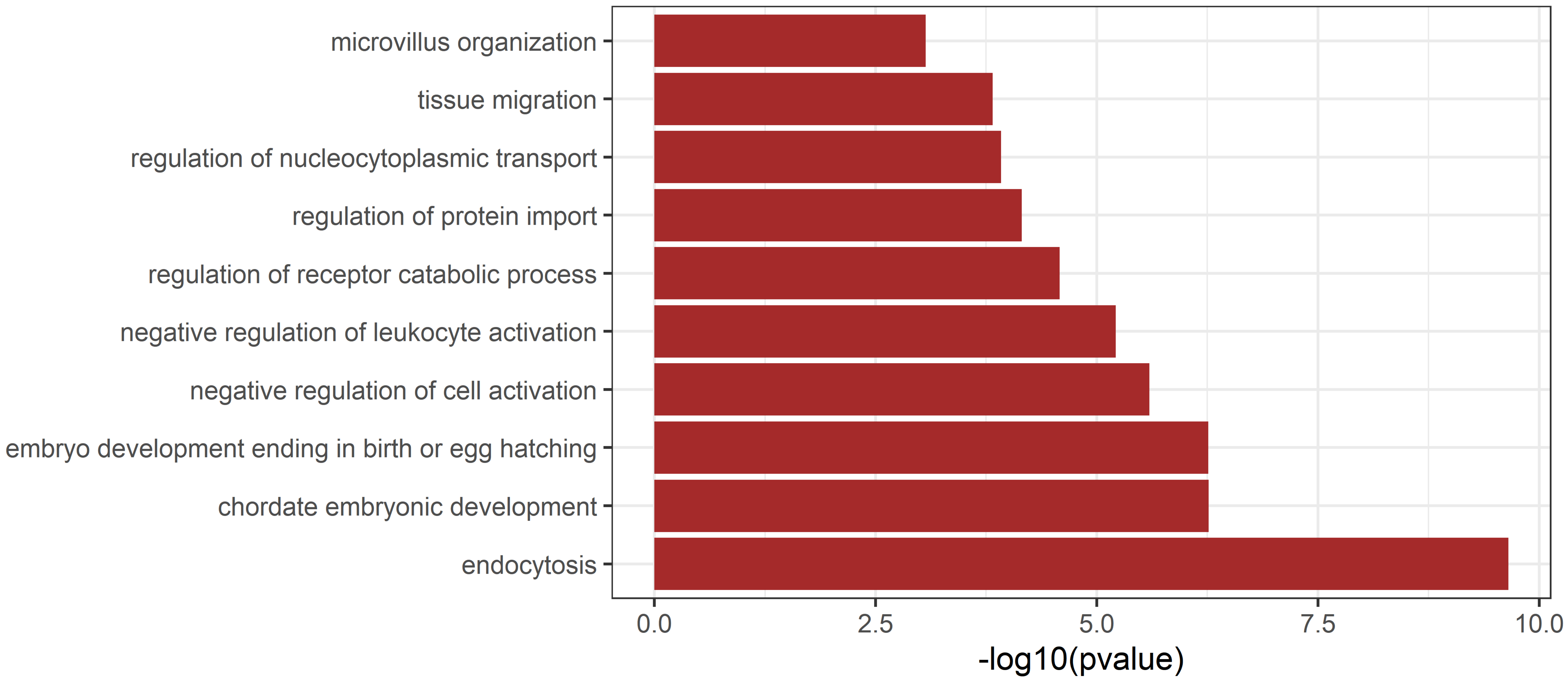


**Fig. S4 GO enrichment for MER41-enhancers annotated in human placenta**
The bar plot visualizes the representative GO terms enriched for MER41-enhancers annotated in human placenta. GO enrichment analysis was performed with GREAT using default settings. Only GO terms from the category "Biological Process" are included.

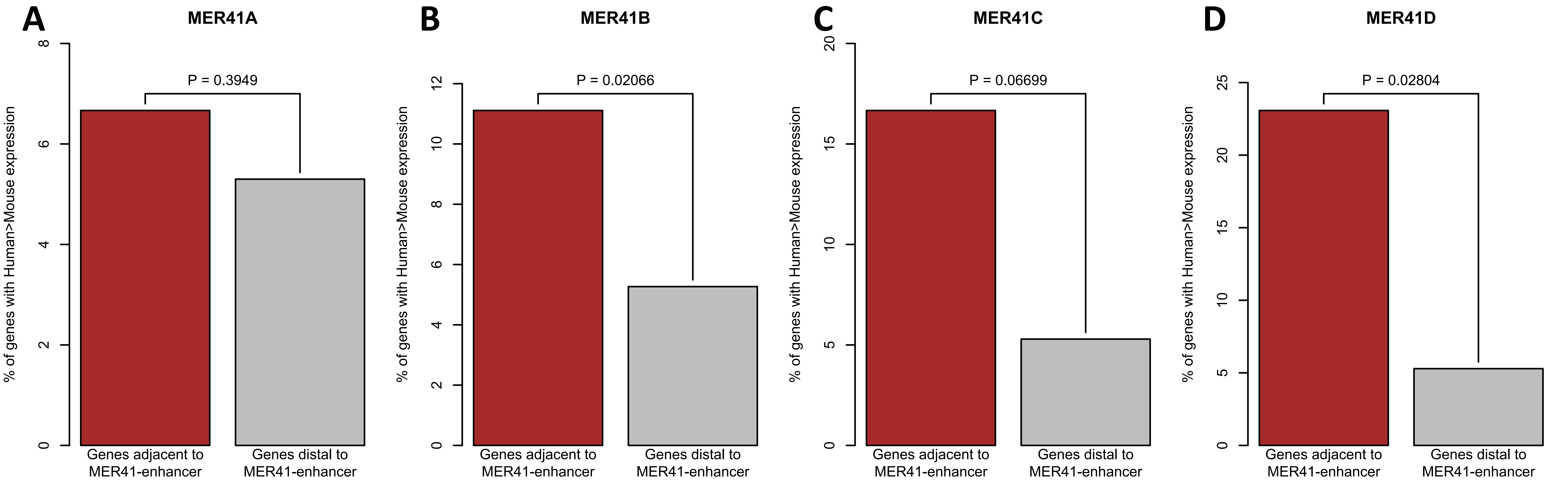


**Fig. S5 Comparison of the genes adjacent or distal to MER41-enhancers regarding their differential expression between human and mouse placentae**

The bar plots compare the proportion of MER41-enhancer adjacent and distal genes that have increased expression in human placenta relative to mouse. MER41-enhancer adjacent genes are defined as those with their TSSs within 50 KB from any MER41-enhancers. P-values calculated using Fisher's Exact Test are indicated. The four MER41 subfamilies (ie. MER41A/B/C/D), which have at least adjacent genes, are included for analysis. MER41E/G are excluded because their adjacent genes are of too small numbers (4 and 0, respectively). This figure is related to Fig. 1B.

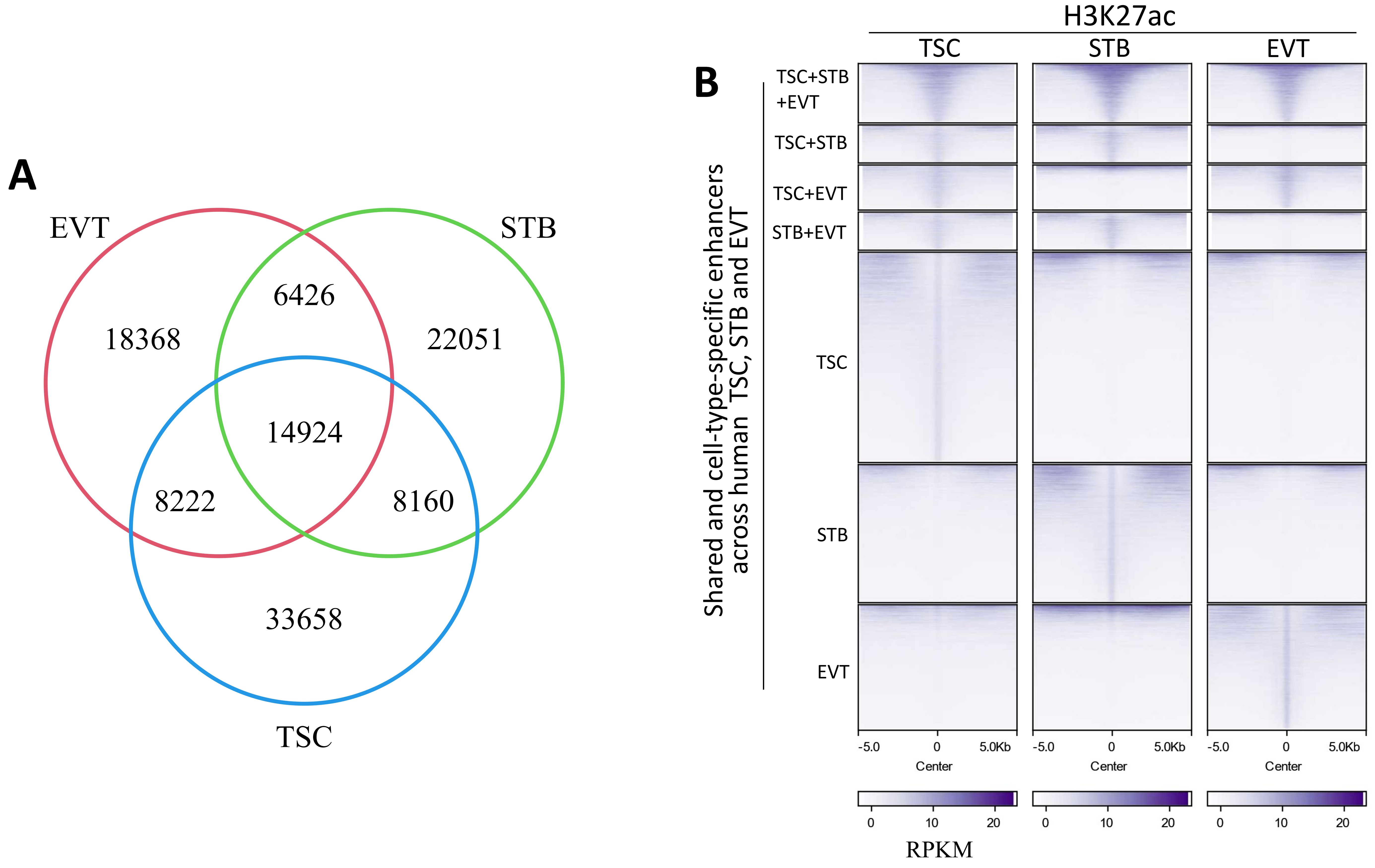


**Fig. S6 Comparison of the epigenetically-annotated enhancers for different types of human trophoblast cells**

**A**.Venn diagram shows the overlapping of the enhancers annotated for human TSC, STB and EVT samples. **B**. Heatmaps show the epigenomic profiles for the enhancers shared or cell-specific for different types of human trophoblast cells. The color gradients indicate the RPKM values (with Input subtracted) calculated from the H3K27ac ChIP-Seq data.

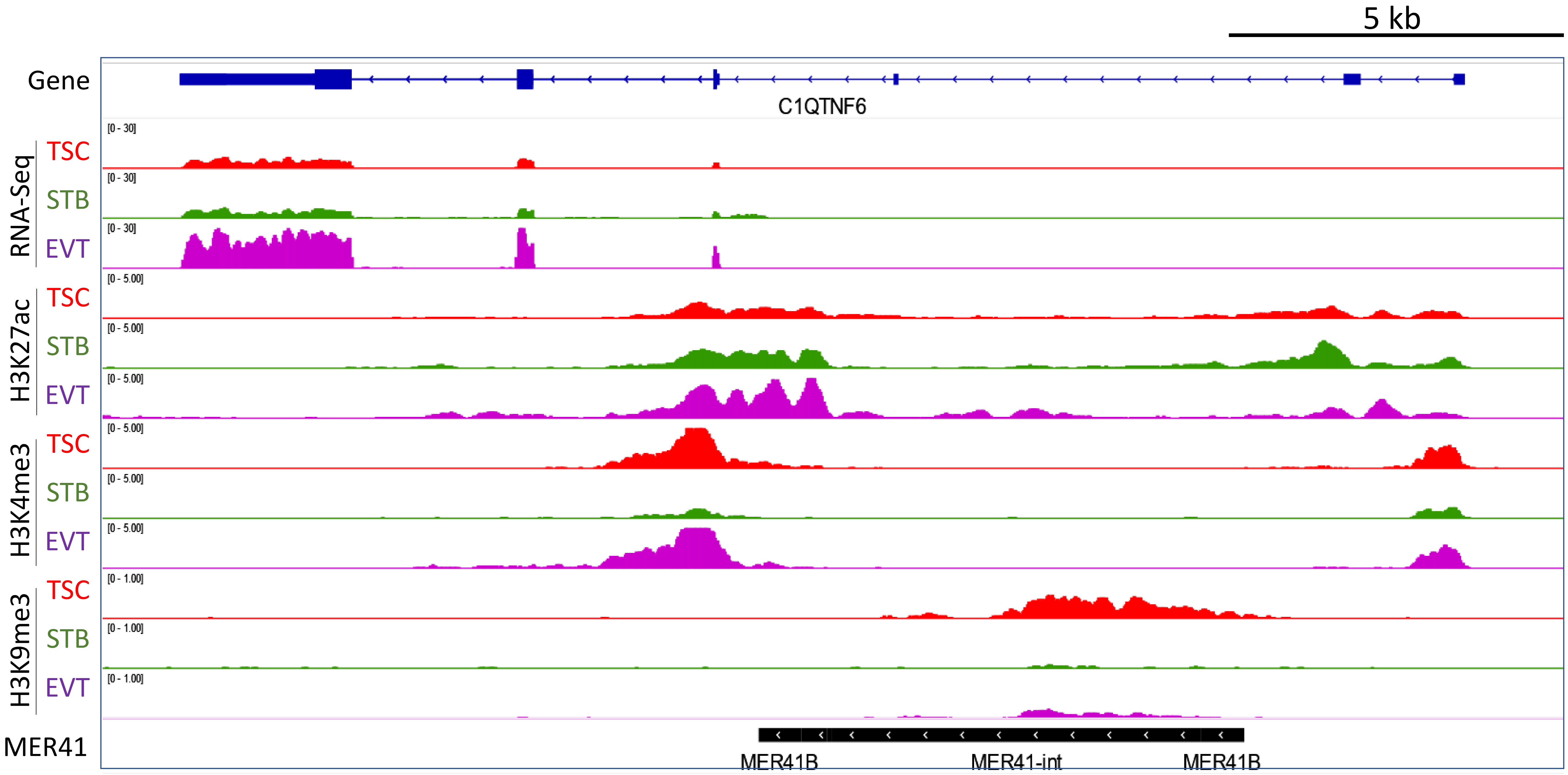


**Fig. S7 IGV tracks showing the transcriptomic and epigenomic profiles surrounding the representative MER41-enhancer associated with *C1QTNF6* gene**

This IGV track is related to Fig. 1F, which shows the patterns for a representative MER41-enhancer adjacent to another gene named *FBN2*.

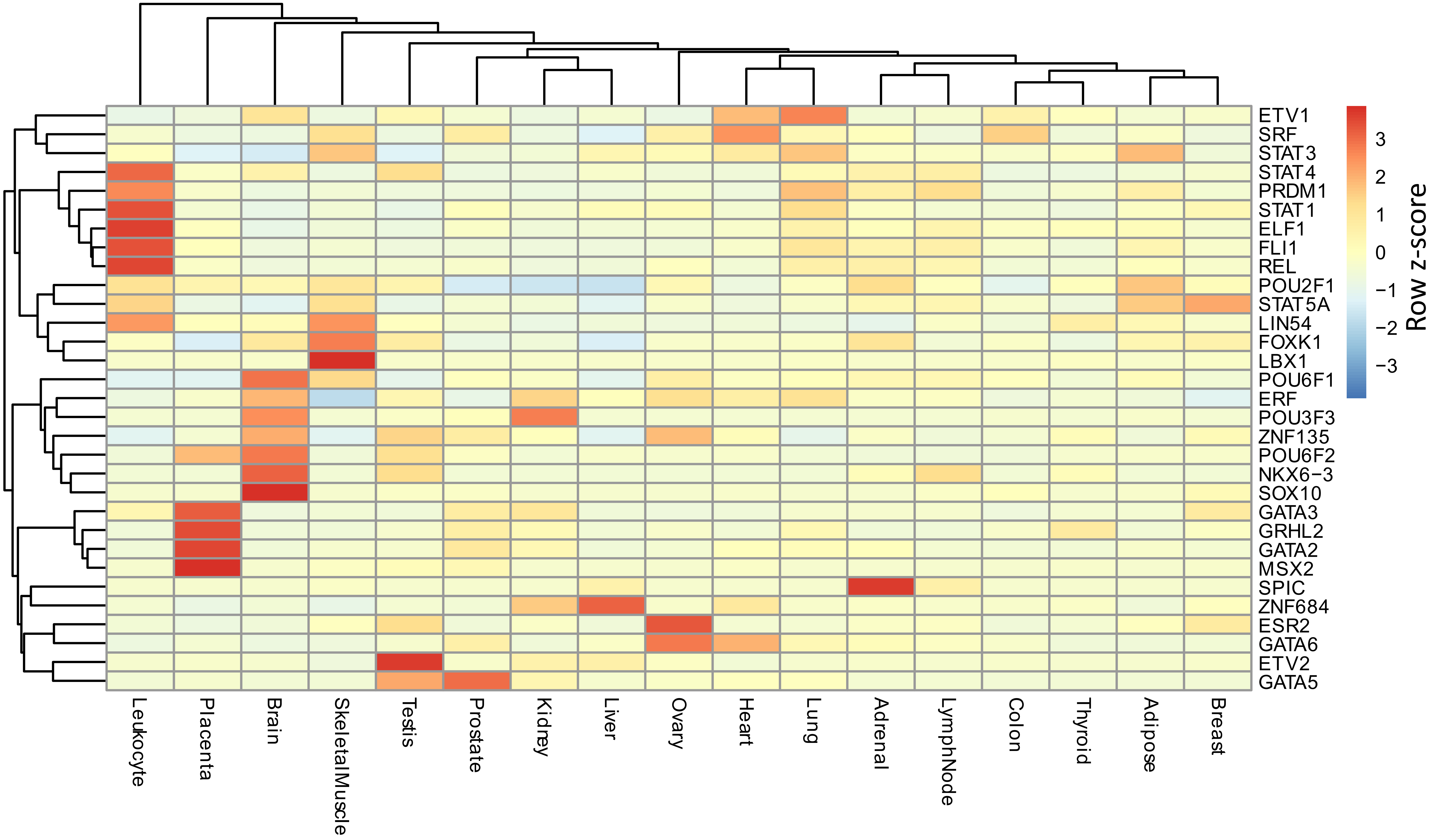


**Fig. S8 Expression profiles for the TFs with their motifs harbored in MER41B sequence**

This heatmap demonstrates the expression profiles for all the TFs with their binding motifs present within MER41B sequence. Four TFs, including GATA2/3, MSX2 and GRHL2, have placenta-enriched expression. The color gradients represent the row z-score calculated from the normalized TPM values.

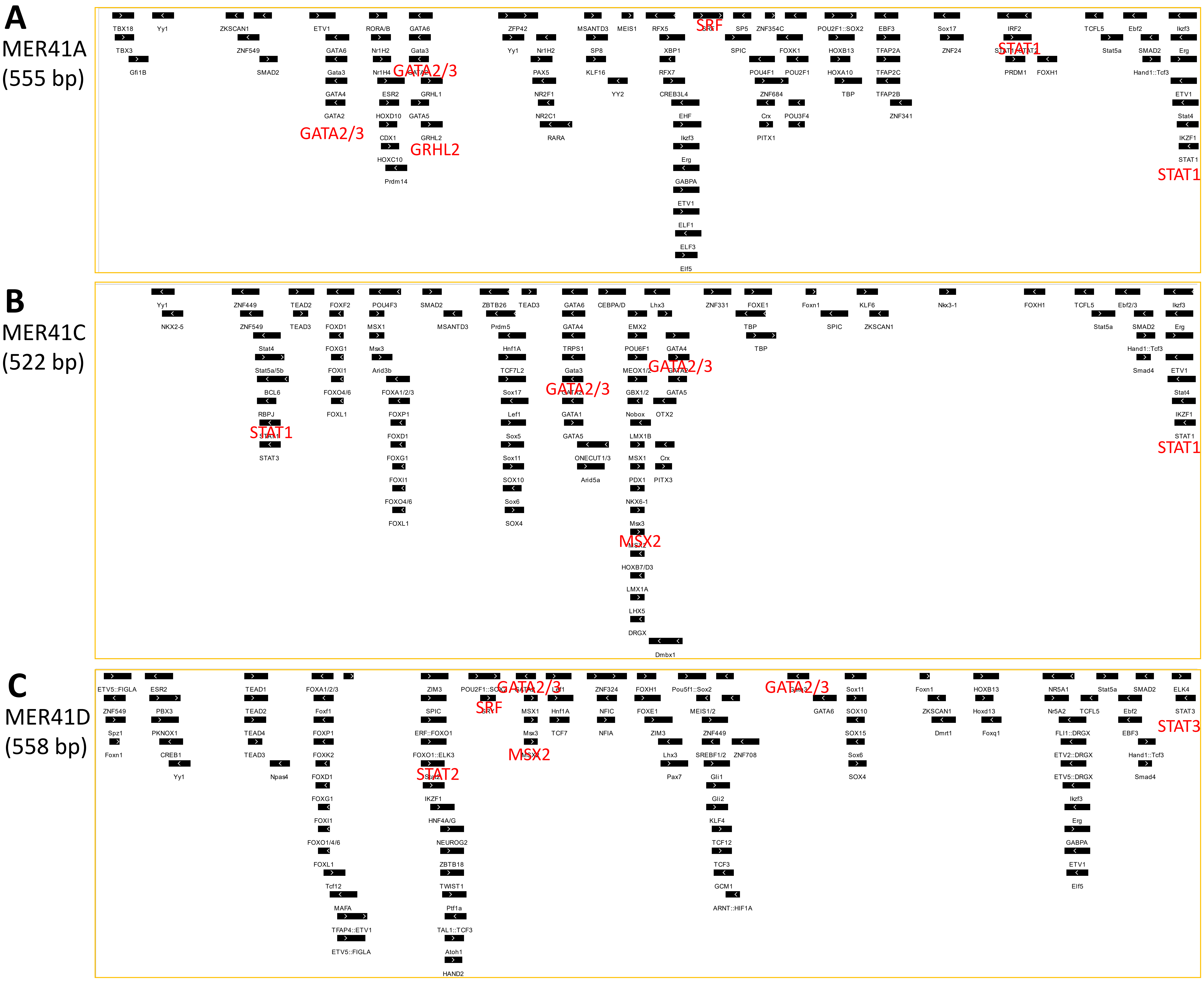


**Fig. S9 Motif configurations of the consensus sequences for MER41A/C/D**

Here shows the motif configurations for the consensus sequences for MER41A (A), MER41C (B) and MER41D (C), respectively. The motifs for several important TFs (STAT1/2/3, SRF, GATAA2/3, MSX2 and GRHL2) are highlighted in red. This figure is related to Fig. 2A.

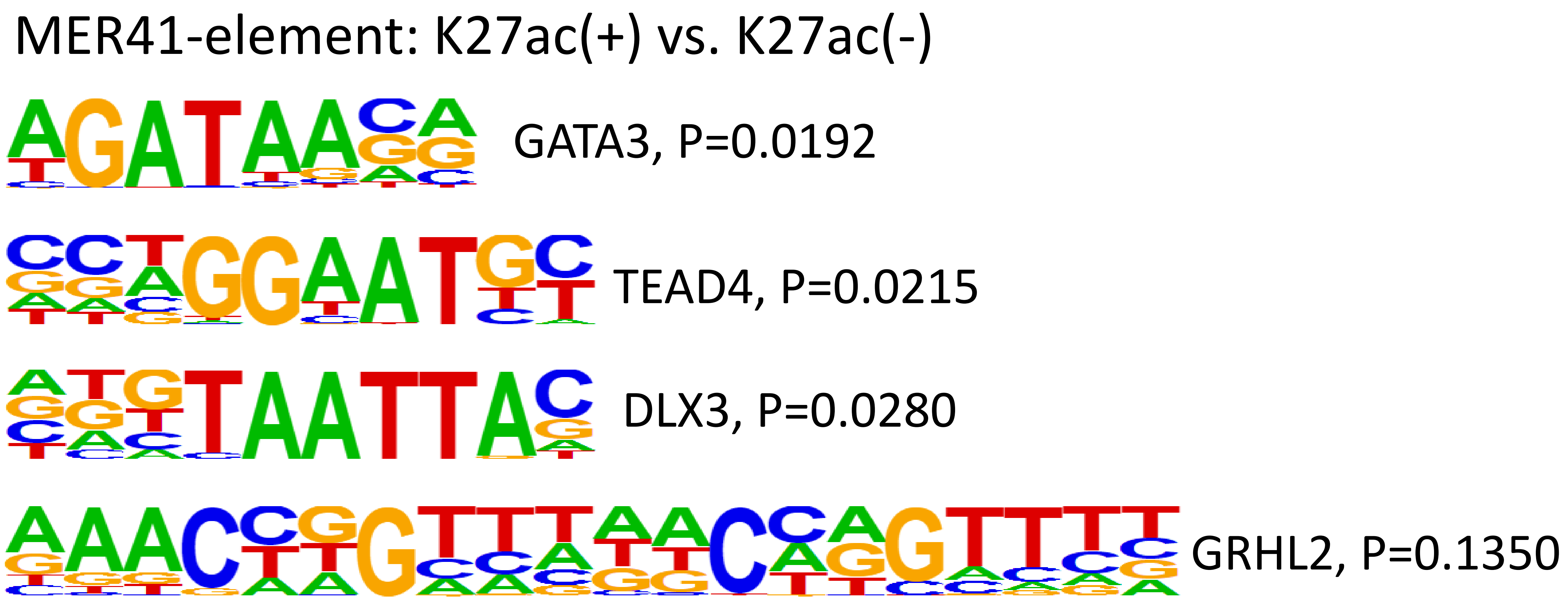


**Fig. S10 Enrichment of the motifs for multiple trophoblast TFs in MER41-enhancers**

This figure shows the enrichment of the motifs for several known trophoblast TFs (GATA3, TEAD4, DLX3, GRHL2) in MER41-enhancers relative to MER41-elements that lack the H3K27ac mark. Motif enrichment analysis was performed using the findMotifs.pl script from the HOMER package. The adjusted p-values are indicated.

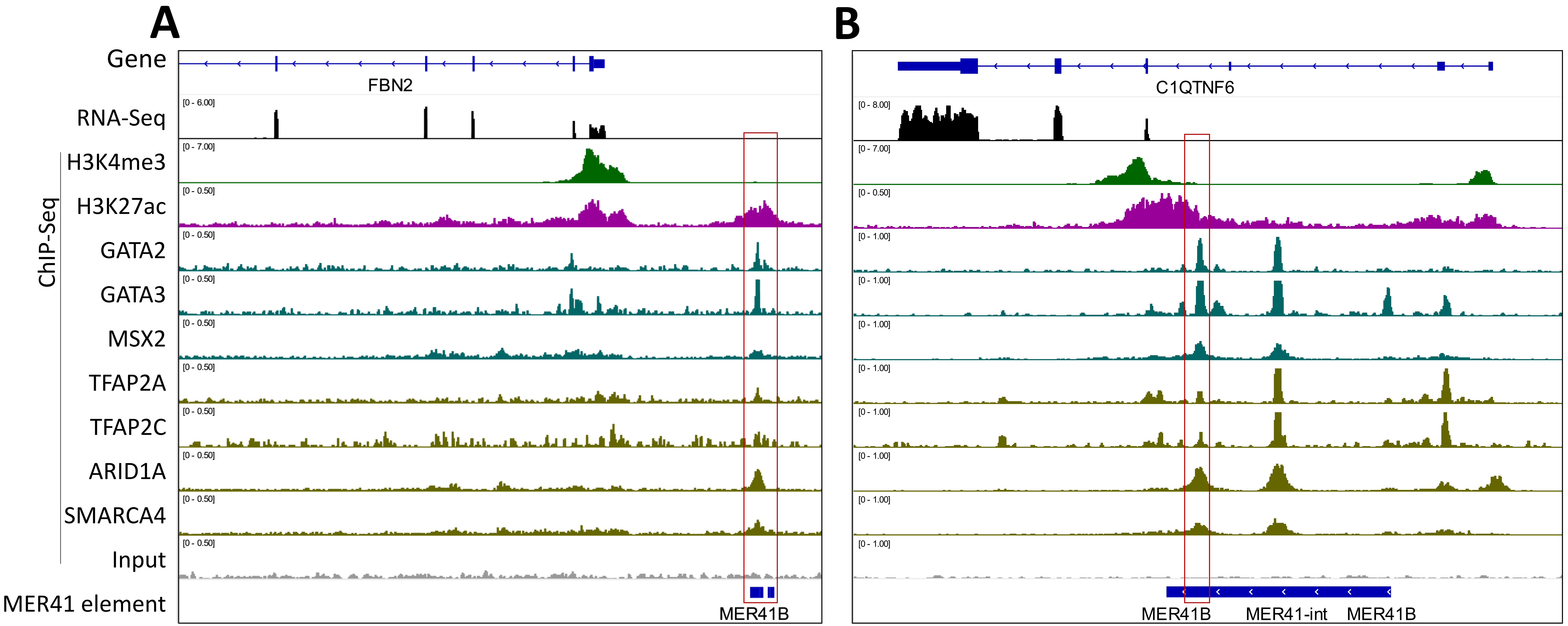


**Fig. S11 Binding of multiple trophoblast TFs and co-factors on representative MER41-enhancers in human TSCs**

This figure shows the binding of GATA2/3, MSX2 and their co-factors on representative MER41-enhancers adjacent to *FBN2* (**A**) and *C1QTNF6* (**B**).

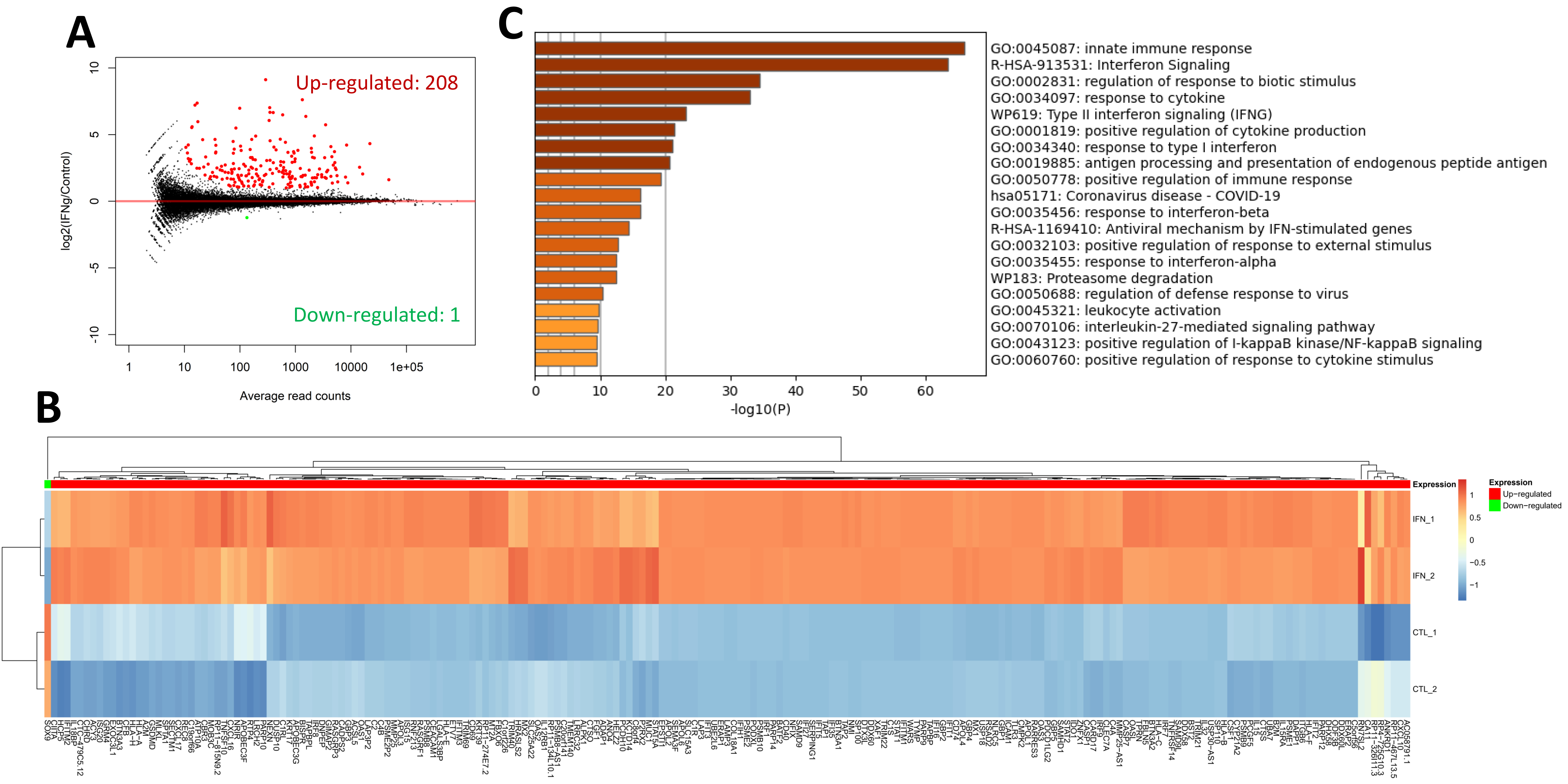


**Fig. S12 Characterization of IFN-stimulated genes in human TSCs**

**A**. MA-plot shows the differentially expressed genes after IFN-stimulation in human TSCs. **B**. Heatmap shows the expression profiles of the differentially expressed genes by IFN-stimulation in human TSCs. **C**. GO enrichment results for IFN-stimulated genes identified in human TSCs.

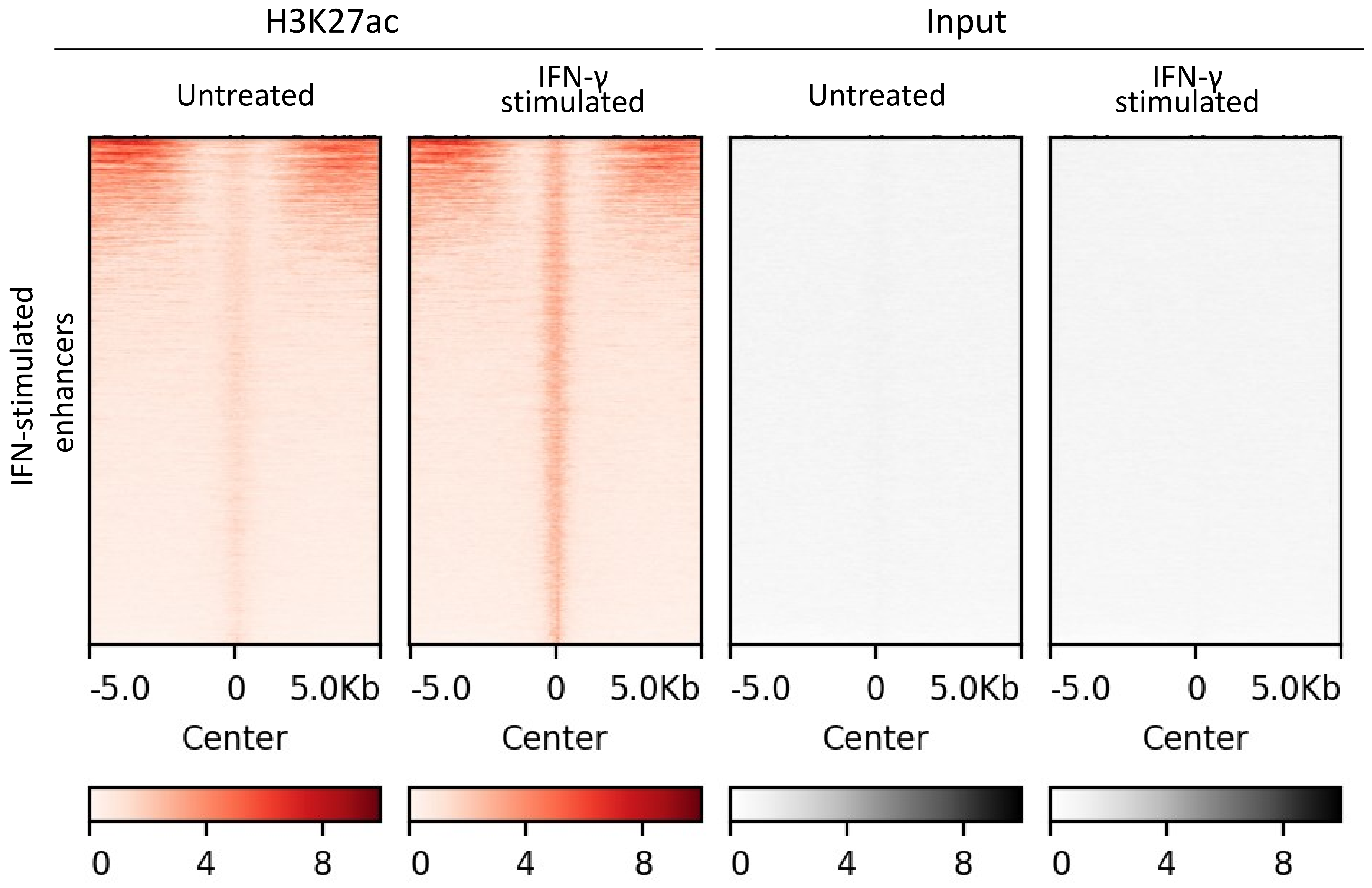


**Fig. S13 Epigenetic profiles for the IFN-stimulated enhancers identified in human TSCs**
The heatmap shows the H3K27ac intensity of IFN-stimulated enhancers identified in human TSCs. The color bar indicates the RPKM values calculated from ChIP-Seq data.

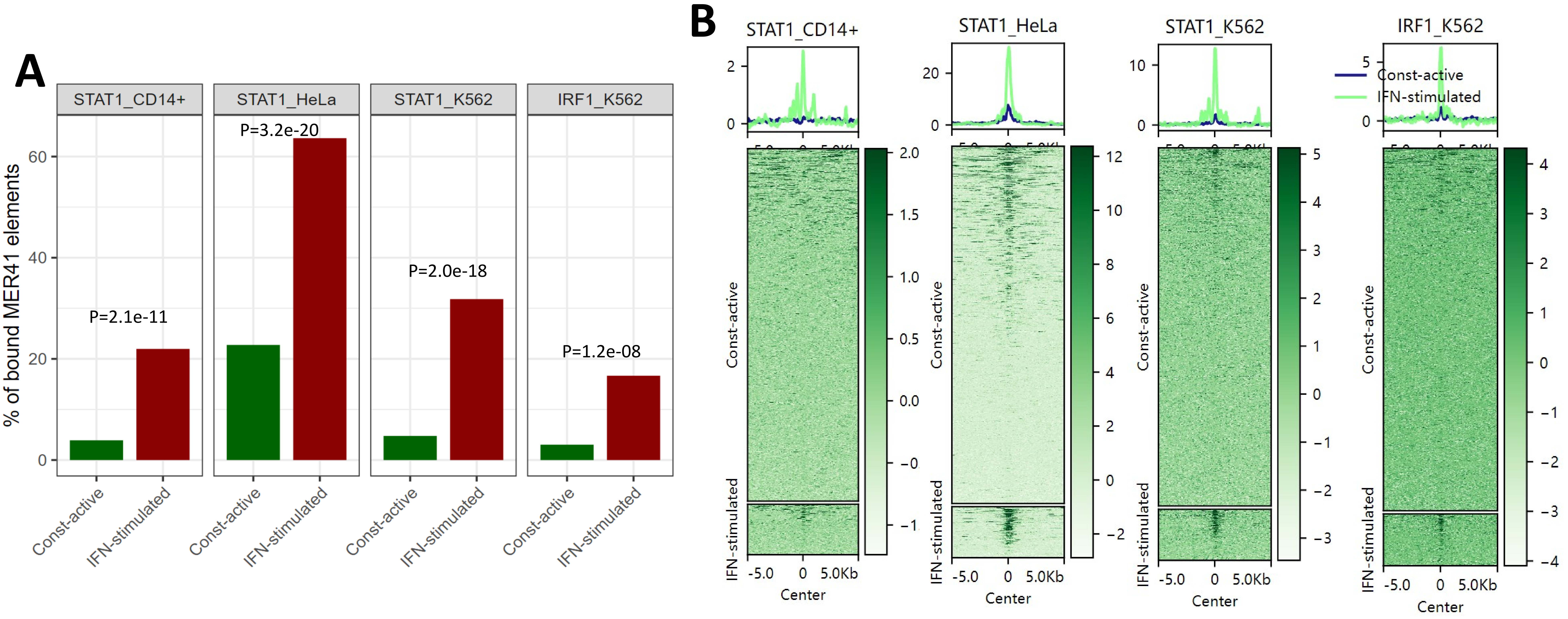


**Fig. S14 Comparison of the binding of immune TFs on the constitutively-active and IFN-stimulated MER41-enhancers annotated in human TSCs**

(A) Bar plots for the binding frequency of immune TFs including STAT1 and IRF1 on the two groups of MER41-enhancers. P-values calculated by using Fisher's Exact Test are indicated. (B) Heatmaps show the binding intensity of STAT1 and IRF1 on the two groups of MER41-enhancers. The color gradients indicate the RPKM values (ChIP – Input) calculated from the ChIP-Seq data. The two groups of MER41-enhancers are first annotated based on the data for human TSCs, and then mapped to the ChIP-Seq data of immune TFs for multiple human cell types for further analysis.

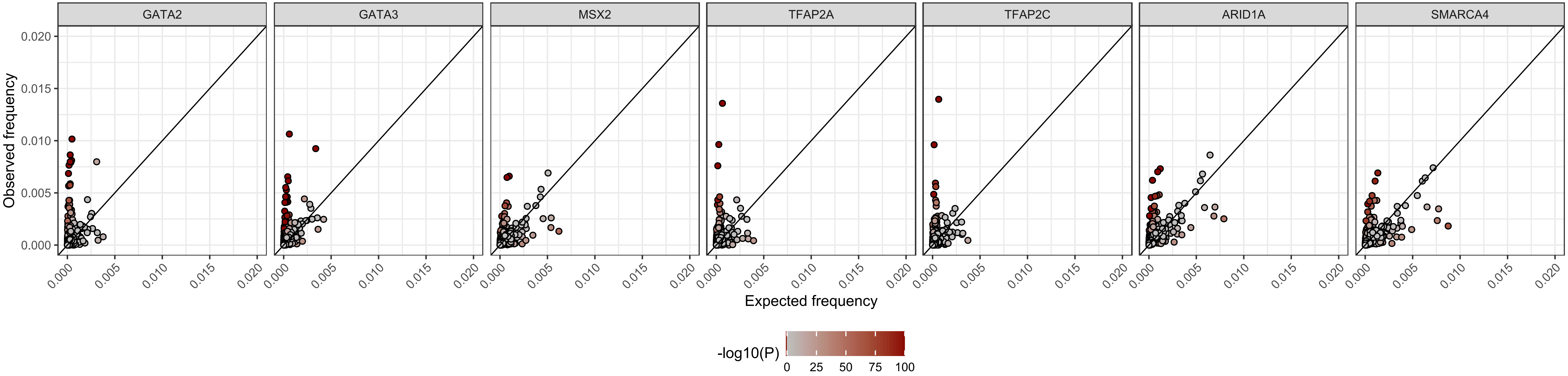


**Fig. S15 Enrichment of specific ERV families within the peaks for different trophoblast TFs**

The scatter plots show the enrichment of different ERV families in the peaks for different trophoblast TFs, including GATA2, GATA3, MSX2 and their co-factors (TFPA2A, TFAP2C, ARID1A and SMARCA4). The color bar indicates the –log10(P).

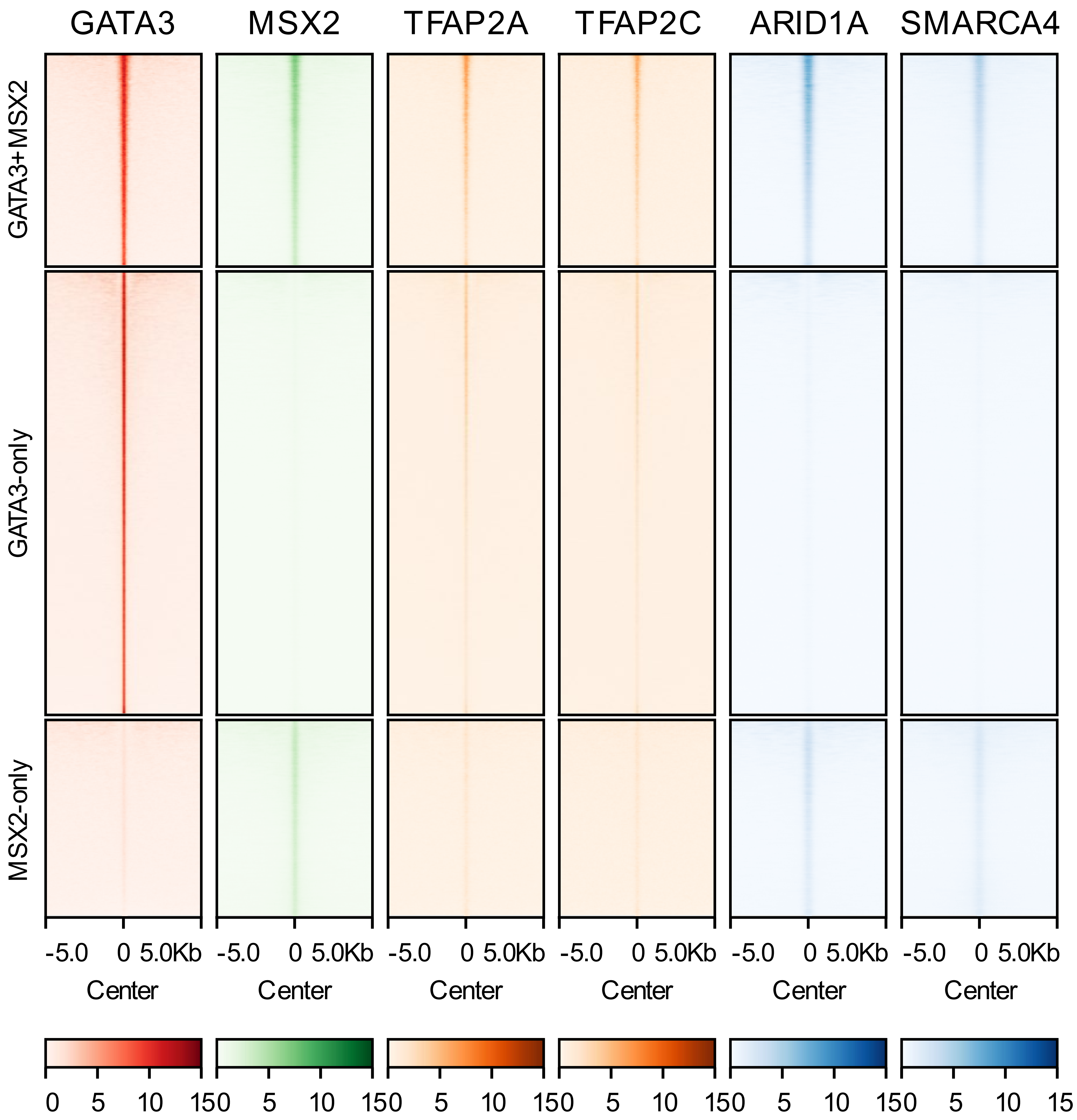


**Fig. S16 Occupancy of H3K27ac and different TFs on GATA3+MSX2, GATA3-only and MSX2-only peaks**
The heatmap shows the ChIP intensity (measured as RPKM value) of GATA3, MSX2 and their related TFs (TFAP2A and TFAP2C for GATA3, and ARID1A and SMARCA4 for MSX2) on peaks bound by GATA3+MSX2, GATA3-only or MSX2-only.

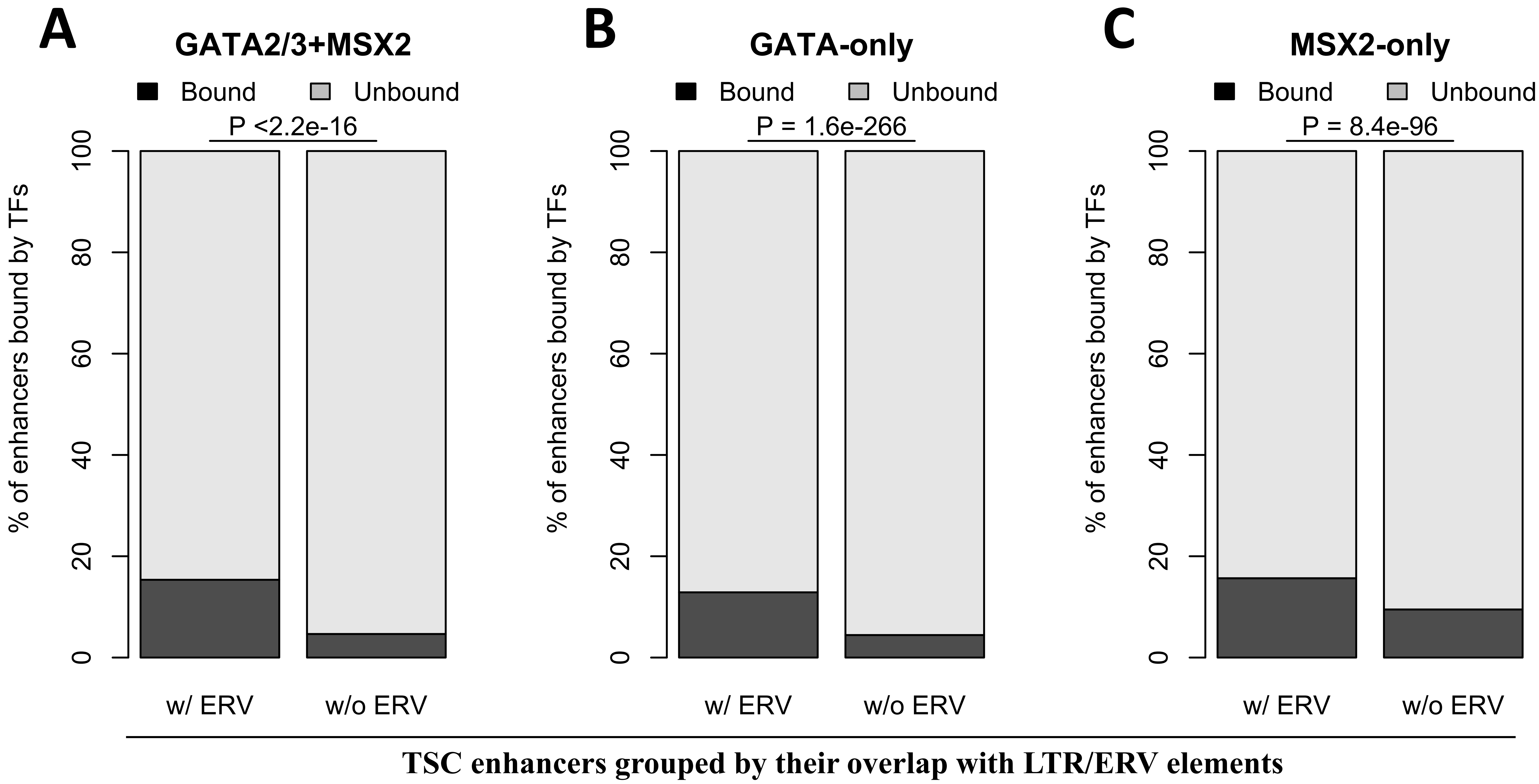


**Fig. S17 Comparison of the percentages of TSC enhancers that are bound by GATA2/3 and MSX2**
The figures compared the percentages of TSC enhancers (ERV-derived or not) that are bound by GATA2/3+MSX2 (A), GATA2/3-only (B) or MSX2-only (C). The peaks of GATA2 and GATA3 were pooled for analysis. P-values calculated by Fisher's Exact Test are denoted.

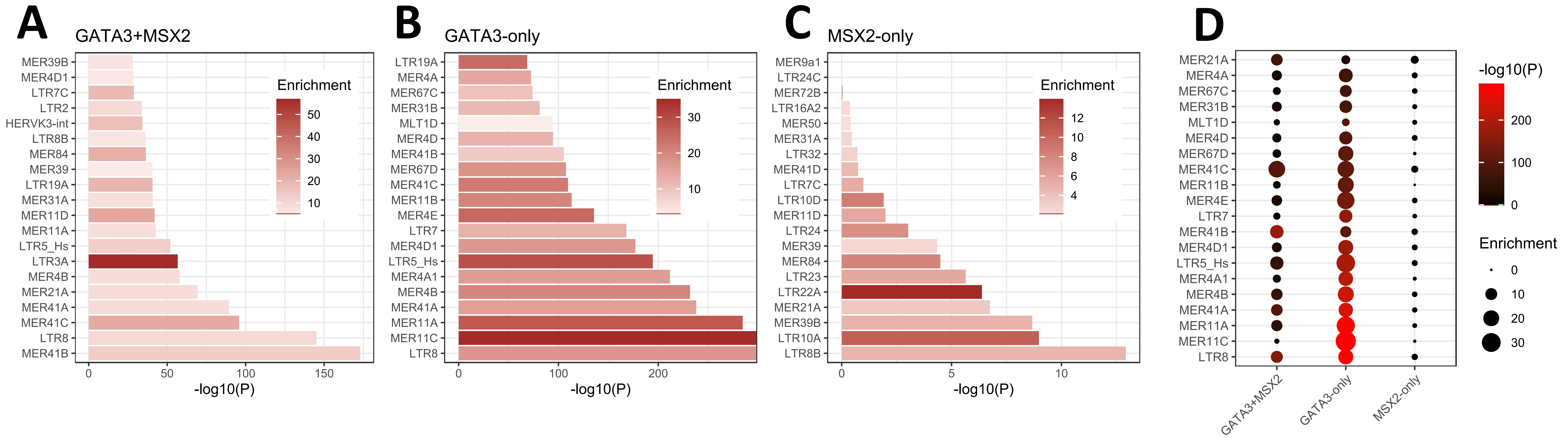


**Fig. S18 Comparison of the enriched ERV families in regions bound by GATA3+MSX2, GATA3-only and MSX2-only**
(A-C) Bar plots show the top 20 enriched ERV families in the regions bound by GATA3+MSX2, GATA3-only and MSX2-only, respectively. The color bar indicates the enrichment fold. The ERV families were selected based on the ERV enrichment result for each group of peaks. (D) Dot plot for the direct comparison of the top 20 enriched ERV families regarding their enrichment in GATA3+MSX2, GATA3-only or MSX2-only peaks. The ERV families were selected after pooling the enrichment results for all groups of peaks.

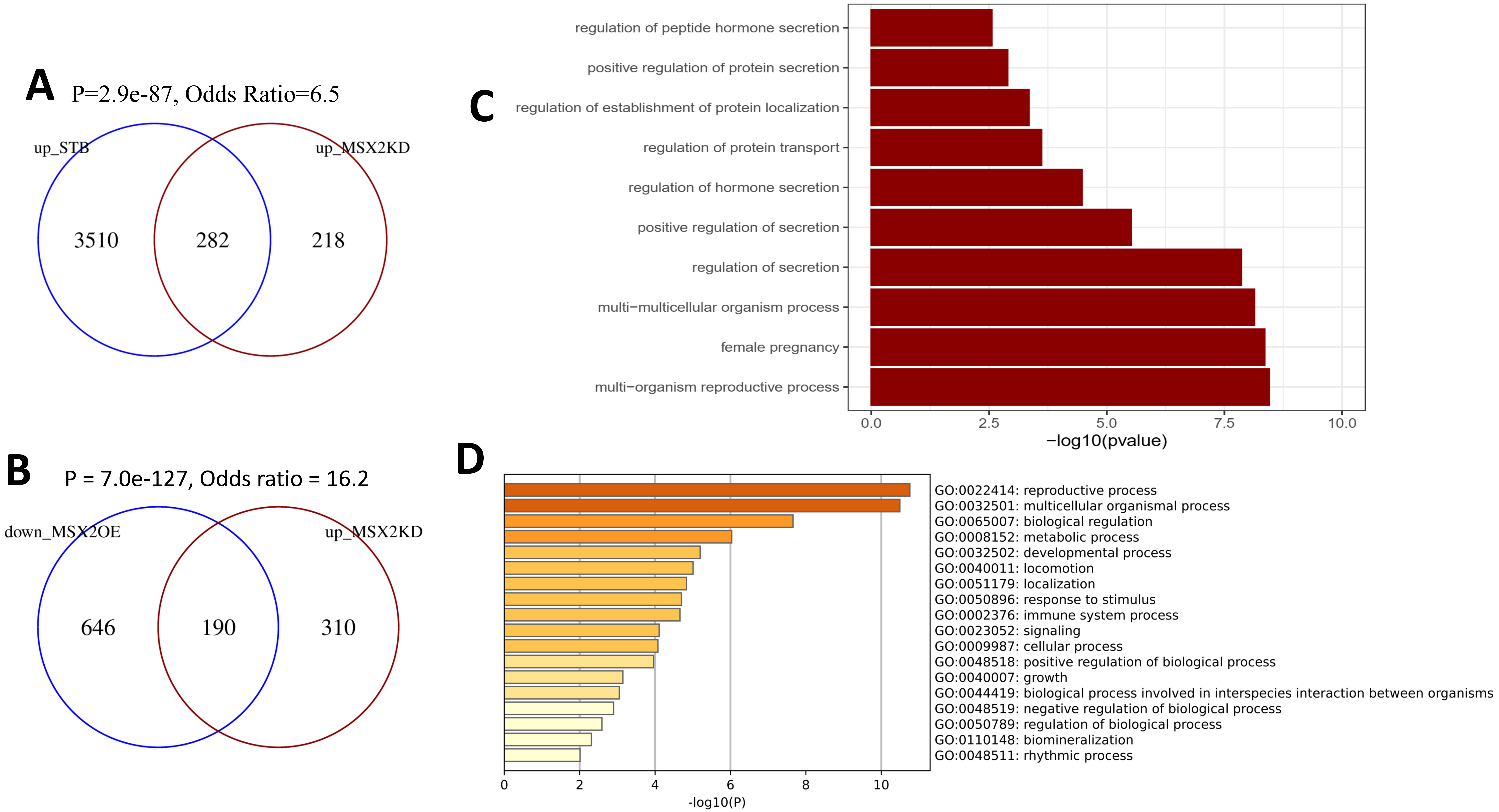


**Fig. S19 Genes under the repression of MSX2 are highly related to placenta function and STB genes**
(**A**) The venn diagram shows the overlap of MSX2-repressed genes and STB-genes. STB genes were identified as genes with significantly higher expression in human STB relative to TSC, through re-analysis of the RNA-Seq data from Okae et al., Cell Stem Cell, 2018. P-value calculated with Fisher's Exact Test was indicated. (**B**) Venn diagram shows the significant overlap of the genes up-regulated in MSX2KD and down-regulated in MSX2OE. The P-value calculated by using Fisher's Exact Test is indicated. (**C**) GO enrichment analysis for the genes that are significantly upregulated in MSX2KD relative to wild-type human TSCs. Top ten GO terms from "Biological process" category were included for visualization. (**D**) GO enrichment result for the 190 intersect genes from B. Only top GO terms from the "Biological Process" category were included for visualization.

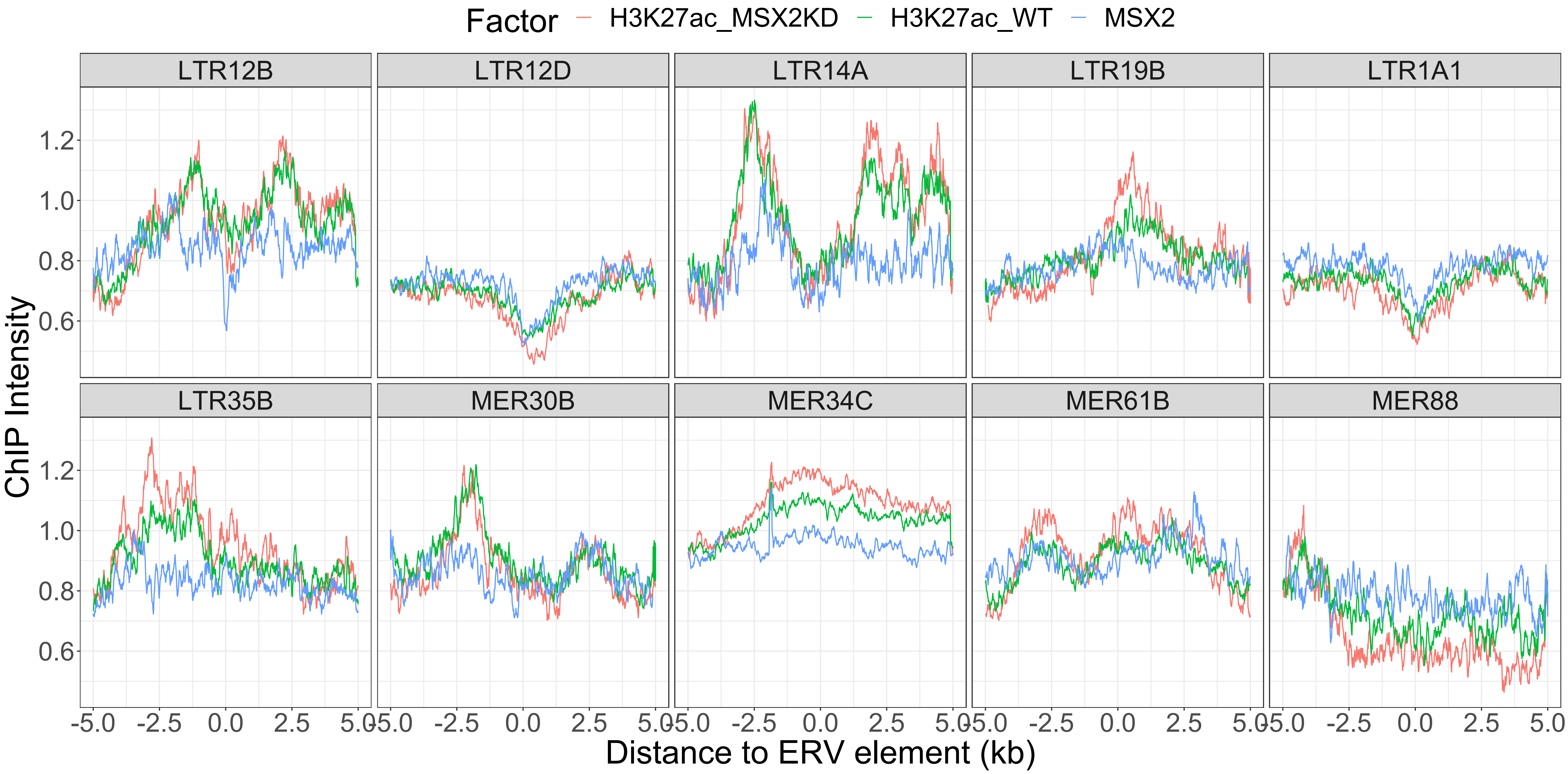


**Fig. S20 H3K27ac levels in WT and MSX2KD human TSCs flanking ERV families unbound by MSX2**
The curves compared the H3K27ac levels in WT and MSX2KD human TSCs flanking ten representative ERV families that are unbound by MSX2. The ChIP intensity was calculated as the FPKM values from the ChIP-Seq data.

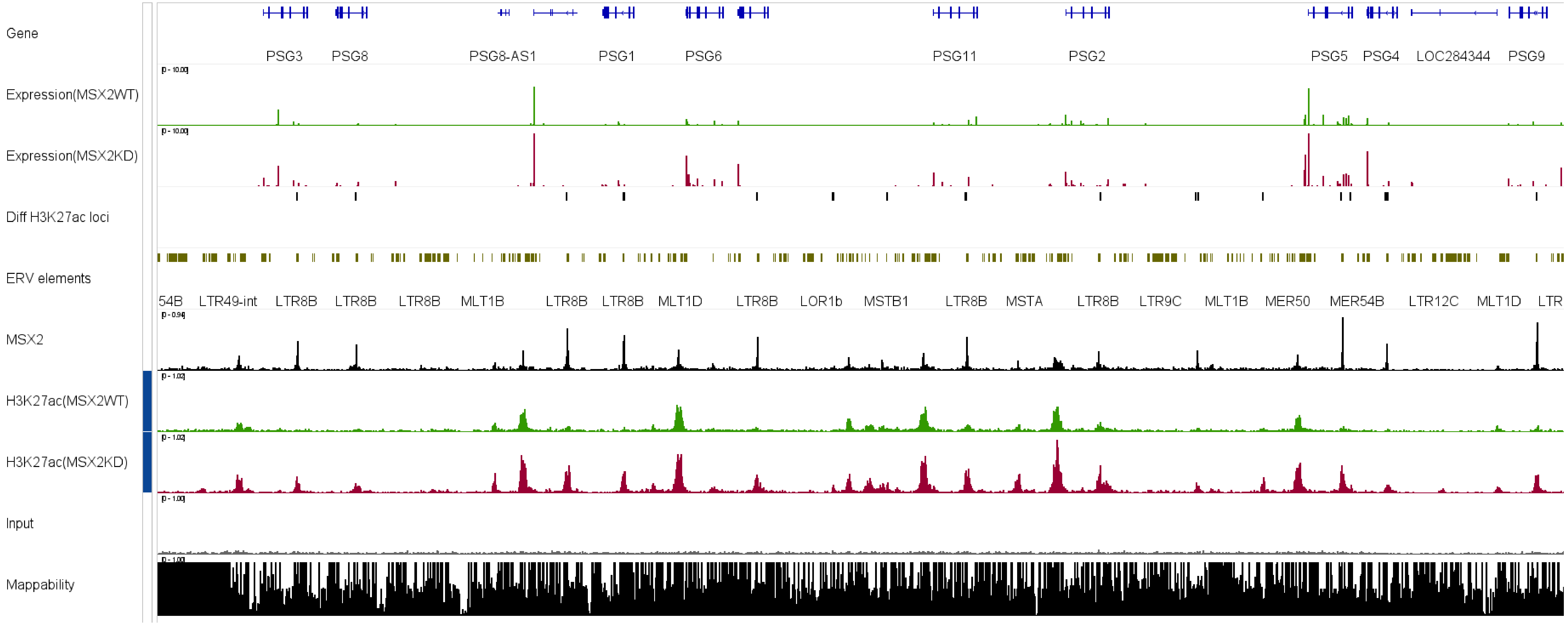


**Fig. S21 Binding of MSX2 on LTR8B-derived cis-elements in multiple PSG genes**

The IGV track shows the transcriptomic and epigenomic (H3K27ac) profiles for the genomic region containing multiple PSG genes. Additional information including differential H3K27ac calling, ERV annotation, MSX2 binding and mappability is also provided side by side.
